# Efficient coalescent simulation and genealogical analysis for large sample sizes

**DOI:** 10.1101/033118

**Authors:** Jerome Kelleher, Alison M. Etheridge, Gil McVean

## Abstract

A central challenge in the analysis of genetic variation is to provide realistic genome simulation across millions of samples. Present day coalescent simulations do not scale well, or use approximations that fail to capture important long-range linkage properties. Analysing the results of simulations also presents a substantial challenge, as current methods to store genealogies consume a great deal of space, are slow to parse and do not take advantage of shared structure in correlated trees. We solve these problems by introducing sparse trees and coalescence records as the key units of genealogical analysis. Using these tools, exact simulation of the coalescent with recombination for chromosome-sized regions over hundreds of thousands of samples is possible, and substantially faster than present-day approximate methods. We can also analyse the results orders of magnitude more quickly than with existing methods.

## 1 Introduction

The coalescent process (Kingman, 1982; Hudson, 1983a) underlies much of modern population genetics and is fundamental to our understanding of molecular evolution. The coalescent describes the ancestry of a sample of *n* genes in the absence of recombination, selection, population structure and other complicating factors. The model has proved to be highly extensible, and these and many other complexities required to model real populations have successfully been incorporated (Wakeley, 2008). Simulation has played a key role in coalescent theory since its beginnings (Hudson, 1983a), partly due to the ease with which it can be simulated: for a sample of *n* genes, we require only *O*(*n*) time and space to simulate a genealogy (Hudson, 1990).

Soon after the single locus coalescent was derived, Hudson defined an algorithm to simulate the coalescent with recombination (Hudson, 1983b). However, after some early successes in characterising this process (Hudson and Kaplan, 1985; Kaplan and Hudson, 1985) little progress was made because of the complex distribution of blocks of ancestral material among ancestors. Some years after Hudson’s pioneering work, the study of recombination in the coalescent was recast in the framework of the Ancestral Recombination Graph (Griffiths, 1991; Griffiths and Marjoram, 1997). In the ARG, nodes are events (either recombination or common ancestor) and the edges are ancestral chromosomes. A recombination event results in a single ancestral chromosome splitting into two chromosomes, and a common ancestor event results in two chromosomes merging into a common ancestor. Analytically, the ARG is a considerable simplification of Hudson’s earlier work as it models all recombination events that occurred in the history of a sample and not just those that can potentially affect the genealogies. Many important results have been derived using this framework, one of which is particularly significant for our purposes here. Ethier and Griffiths (1990) proved that the expected number of recombination events back to the Grand MRCA of a sample of *n* individuals grows like *e^ρ^* as *ρ* →∞, where *ρ* is the population scaled recombination rate. In this paper we consider a diploid model in which we have a sequence of *m* discrete sites that are indexed from zero. Recombination occurs between adjacent sites at rate *r* per generation, and therefore *ρ* = *4N_e_r*(*m −* 1). The Ethier and Griffiths result implies that the time required to simulate an ARG grows exponentially with the sequence length, and we can only ever hope to simulate ARGs for the shortest of sequences.

This result, coupled with the observed poor scaling of coalescent simulators such as the seminal ms program (Hudson, 2002) seems to imply that simulating the coalescent with recombination over chromosome scales is hopeless, and researchers have therefore sought alternatives. The sequentially Markov coalescent (SMC) approximation (McVean and Cardin, 2005; Marjoram and Wall, 2006) underlies the majority of present day genome scale simulation (Chen et al., 2009; Excoffier and Foll, 2011; Staab et al., 2014) and inference methods (Li and Durbin, 2011; Schiffels and Durbin, 2014; Rasmussen et al., 2014). The SMC simplifies the process of simulating genealogies by assuming that each marginal tree depends only on its immediate predecessor as we move from left-to-right across the sequence. As a consequence, the time required to simulate genealogies scales linearly with increasing sequence length. In practice, SMC based simulators such as `MaCS` and `scrm` are many times faster than `ms`.

The SMC has disadvantages, however. Firstly, the SMC discards all long range linkage information and therefore can be a poor approximation when modelling features such as the length of admixture blocks (Liang and Nielsen, 2014). Improving the accuracy of the SMC is also difficult and error prone. For example, the `MaCS` simulator (Chen et al., 2009) has a parameter to increase the number of previous trees on which a marginal tree can depend. Paradoxically, this gives a worse approximation to the coalescent with recombination if we increase it beyond a certain limit (Staab et al., 2014). Incorporating complexities such as population structure (Eriksson et al., 2009), intra-codon recombination (Arenas and Posada, 2010) and inversions (Peischl et al., 2013) is non-trivial and can be substantially more complex than the corresponding modification to the exact coalescent model. Also, while SMC based methods scale well in terms of increasing sequence length, currently available simulators do not scale well in terms of sample size.

We solve these problems by introducing sparse trees and coalescence records as the fundamental units of genealogical analysis. By creating a concrete formalisation of the genealogies generated by the coalescent process in terms of an integer vector, we greatly increase the efficiency of simulating the exact coalescent with recombination. In Section 2 we discuss how Hudson’s classical simulation algorithm can be defined in terms of these sparse trees, and why this leads to substantial gains in terms of the simulation speed and memory usage. We show that our implementation of the exact coalescent, `msprime`, is competitive with approximate simulators for small sample sizes, and is faster than all other simulators for large sample sizes. This is possible because Hudson’s algorithm does not traverse the entire ARG, but rather a small and poorly understood subset of it. We show some preliminary results indicating that the number of nodes in this graph may be a quadratic function of the scaled recombination rate *ρ* rather than an exponential.

Generating simulated data is of little use if the results cannot be processed in an efficient and convenient manner. Currently available methods for storing and processing genealogies perform very poorly on trees with hundreds of thousands of nodes. In Section 3 we show how the encoding of the correlated trees output by the simulation in Section 2 leads to an extremely compact method of storing these genealogies. For large simulations, the representation can be thousands of times smaller than the most compact tree serialisation format currently available. Our encoding also leads to very efficient tree processing algorithms; for example, sequential access to trees is several orders of magnitude faster than existing methods.

The advantages of faster and more accurate simulation over huge sample sizes, and the ability to quickly process very large result sets may enable applications that were not previously feasible. In Section 4 we conclude by discussing some of these applications and other uses of our novel encoding of genealogies. The methods developed in this paper allow us to simulate the coalescent for very large sample sizes, where the underlying assumptions of the model may be violated (Wakeley and Takahashi, 2003; Bhaskar et al., 2014). Addressing these issues is beyond the scope of this work, but we note that the majority of our results can be applied to simulations of any retrospective population model.

## 2 Efficient coalescent simulation

In this section we define our encoding of coalescent genealogies, and show how this leads to very efficient simulations. There are many different simulation packages, and so we begin with a brief review of the state-of-the-art before defining our encoding in Section 2.1 and analysing its performance in Section 2.2.

Two basic approaches exist to simulate the coalescent with recombination. The first approach was defined by Hudson (1983b), and works by applying the effects of recombination and common ancestor events to the ancestors of the sample as we go backwards in time. Events occur at a rate that depends only on the state of the extant ancestors, and so we can generate the waiting times to these events efficiently without considering the intervening generations. This contrasts with time-reversed generation-by-generation methods (Excoffier et al., 2000; Laval and Excoffier, 2004; Anderson et al., 2005; Liang et al., 2007) which are more flexible but also considerably less efficient. The first simulation program published based on Hudson’s algorithm was `ms` (Hudson, 2002). After this, many programs were published to simulate various evolutionary complexities not handled by `ms`, such as selection (Spencer and Coop, 2004; Teshima and Innan, 2009; Ewing and Hermisson, 2010; Shlyakhter et al., 2014), recombination hotspots (Hellenthal and Stephens, 2007), codon models (Arenas and Posada, 2007), intra-codon recombination (Arenas and Posada, 2010) and models of species with a skewed offspring distribution (Zhu et al., 2015). Others developed user interfaces to facilitate easier analysis (Mailund et al., 2005; Ramos-Onsins and Mitchell-Olds, 2007).

The second fundamental method of simulating the coalescent with recombination is due to Wiuf and Hein (1999a). In Wiuf and Hein’s algorithm we begin by generating a coalescent tree for the left-most locus and then move across the sequence, updating the genealogy to account for recombination events. This process is considerably more complex than Hudson’s algorithm because the relationship between trees as we move across the genome is non-Markovian: each tree depends on all previously generated trees. Because of this complexity, exact simulators based on Wiuf and Hein’s algorithm are significantly less efficient than `ms` (Staab et al., 2014; Wang et al., 2014). However, Wiuf and Hein’s algorithm has provided the basis for the SMC approximation (McVean and Cardin, 2005; Marjoram and Wall, 2006), and programs based on this approach (Chen et al., 2009; Excoffier and Foll, 2011; Staab et al., 2014) can simulate long sequences far more efficiently than exact methods such as `ms`. Very roughly, we can think of Wiuf and Hein’s algorithm performing a depth-first traversal of the ARG, and Hudson’s algorithm a breadth-first traversal. Neither explore the full ARG, but instead traverse the subset required to contruct all marginal genealogies.

Recently, Hudson’s algorithm has been utilised in `cosi2` (Shlyakhter et al., 2014), which takes a novel approach to simulating sequences under the coalescent. The majority of simulators first generate genealogies and then throw down mutations in a separate process. In `cosi2` these two processes are merged, so that mutations are generated during traversal of the ARG. Instead of associating a partial genealogy with each ancestral segment, `cosi2` maps ancestral segments directly to the set of sampled individuals at the leaves of this tree. When a coalescence between two overlapping segments occurs, we then have sufficient information to generate mutations and map them to the affected samples. This strategy, coupled with the use of sophisticated data structures, makes `cosi2` many times faster than competing simulators such as `msms`. The disadvantage of combining the mutation process with ARG traversal, however, is that the underlying genealogies are not available, and `cosi2` cannot directly output coalescent trees.

Many reviews are available to compare the various coalescent simulators in terms of their features (Carvajal-Rodríguez, 2008; Liu et al., 2008; Arenas, 2012; Yuan et al., 2012; Hoban et al., 2012; Yang et al., 2014). Little information is available, however, about their relative efficiencies. Hudson’s `ms` is widely regarded as the most efficient implementation of the exact coalescent and is the benchmark against which other programs are measured (Marjoram and Wall, 2006; Chen et al., 2009; Excoffier and Foll, 2011; Staab et al., 2014; Yang et al., 2014; Wang et al., 2014). However, for larger sample sizes and long sequence lengths, `msms` (Ewing and Hermisson, 2010) is much faster than `ms`. Also, for these larger sequence lengths and sample sizes, `ms` is unreliable and crashes (Excoffier and Foll, 2011; Yang et al., 2014). Thus, `msms` is a much more suitable baseline against which to judge performance. The `scrm` simulator is the most efficient SMC based method currently available (Staab et al., 2014).

### 2.1 Hudson’s algorithm with sparse trees

An oriented tree (Knuth, 2011, p. 461) is a sequence of integers π_1_π_2_ …, such that π*_u_* is the parent of node *u* and *u* is a root if *π_u_* = 0. For example, the trees

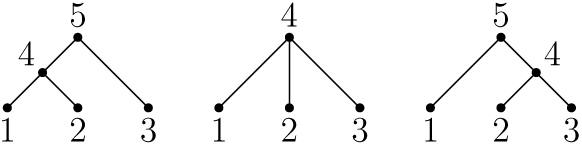

are defined by the sequences 〈4, 4, 5, 5, 0〉, 〈4, 4, 4, 0〉 and 〈5, 4, 4, 5, 0〉, respectively. Oriented trees provide a concise and efficient method of representing genealogies, and have been used in coalescent simulations of a spatial continuum model (Kelleher et al., 2013a, 2014). These simulations adopted the convention that the individuals in the sample (leaf nodes) are mapped to the integers 1,…, *n*. For every internal node *u* we have *n* < *u* < 2*n* and (for a binary tree) the root is 2*n* − 1. We refer to such trees as dense because the 2*n* − 2 non-zero entries of the (binary) tree *π* occur at *u* = 1,…, 2*n* − 2. A sparse oriented tree (or more concisely, sparse tree) is an oriented tree *π* in which the leaf nodes are 1,…, *n* as before, but internal nodes can be any integer > *n*.

Each ancestor in the history of the sample in which at least one marginal coalescence occurred corresponds to exactly one node. Ancestral nodes are numbered sequentially from *n* + 1. Note that we make a distinction between common ancestor events and coalescence events throughout. A common ancestor event occurs when two ancestors merge to form a common ancestor. If these ancestors have overlapping ancestral material, then there will also be at least one coalescence event, which is defined as a single contiguous block of sequence coalescing within a common ancestor. In Hudson’s algorithm there are many common ancestor events that do not result in coalescence, and it is important to distinguish between them.

Let the tuple (*ℓ, r, u*) define a segment carrying ancestral material. This segment represents the mapping of the half-closed genomic interval [*ℓ*, r) to the tree node *u*. Each ancestor a is defined by a set of non-overlapping segments. Initially we have *n* ancestors, each consisting of a single segment (0, *m, u*) for 1 ≤ *u* ≤ *n*. The only other state required by the algorithm is the time *t*, and the next node *w*; initially, *t* = 0 and *w* = *n* +1.

Let *P* be the set of ancestors at a given time *t*. Recombination events happen at rate *ρL/*(*m −* 1) where

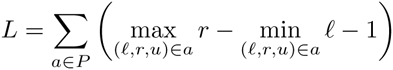
is the number of available ‘links’ that may be broken. (We use a fixed recombination rate here for simplicity, but an arbitrary recombination map can be incorporated without difficulty.) We choose one of the available breakpoints uniformly, and split the ancestry of the individual at that point into two recombinant ancestors. If this breakpoint is at *k*, we assign all segments with *r* ≤ *k* to one ancestor and all segments with *ℓ* ≥ *k* to the other. If there is a segment (*ℓ, r, u*) such that *ℓ* < *k* < *r*, then *k* falls within this segment and it is split such that the segment (*ℓ, k, u*) is assigned to one ancestor and (*k, r, u*) is assigned to the other.

Common ancestor events occur at rate |*P*|(|*P*| − 1). Two ancestors *a* and *b* are chosen and their ancestry merged to form their common ancestor. If their segments do not overlap, the set of ancestral segments of the common ancestor is the union of those of *a* and *b*. If segments do overlap, we have coalescence events which must be recorded. We define a coalescence event as the merging of two segments over the interval [*ℓ*, *r*) into a single ancestral segment. In general the coordinates of overlapping segments *x* and *y* will not exactly coincide, in which case we create an equivalent set of segments by subdividing into the intersections and ‘overhangs’. Suppose then that we have two segments (*ℓ*, *r, u*) and (*ℓ*, *r, v*) from *a* and *b* respectively; over the interval [*ℓ*, *r*) the nodes *u* and *v* coalesce into a common ancestor, which we associate with the next available node *w*. We record this information by storing the coalescence record (*ℓ*, *r, w*, (*u, v*)*, t*). As we see in Section 3.2, these records provide sufficient information to later recover all marginal trees. After recording this coalescence, we then check if there are any other segments in *P* that intersect with [*ℓ*, *r*). If there are, the simulation of this region is not yet complete and we insert the segment (*ℓ*, *r, w*) into the ancestor of *a* and *b*. On the other hand, if there is some subset of [*ℓ*, *r*) which intersects with no other segments in *P*, we know that the marginal tree covering this interval is complete and therefore we do not need to trace its history any further. If any other intervals overlap in *a* and *b*, we perform the same operations, and finally update the next available node by incrementing *w*. In this way, all coalescing intervals within the same ancestor map to the same node w, even if they are disjoint. Conversely, if two disjoint marginal trees contain the same node, we know that this is because multiple segments coalesced simultaneously within the same ancestor.

The algorithm continues generating recombination and common ancestor events at the appropriate rates until *P* is empty, and all marginal trees are complete. This interpretation of Hudson’s algorithm differs from the standard formulations (Hudson, 1983b, 1990; McVean and Cardin, 2005) by concretely defining the representation of ancestry and by introducing the idea of coalescence records. We have omitted many important details here in the interest of brevity; see Appendix A for a detailed listing of our implementation of Hudson’s algorithm, and Appendix B for an illustration of a complete invocation of the algorithm.

There are several advantages to our sparse tree representation of ancestry. Firstly, we do not need to store partially built trees in memory, and the only state we need to maintain is the set of ancestral segments. This leads to substantial time and memory savings, since we no longer have to copy partially built trees at recombination events or update them during coalescences. We can also actively defragment the segments in memory. For example, suppose that as a result of a common ancestor event we have two segments (*ℓ, k, u*) and (*k, r, u*) in an ancestor. We can replace these segments with the equivalent segment (*ℓ,r,u*). Such defragmentation yields significant time and memory savings.

We have developed an implementation of Hudson’s algorithm called `msprime` based on these ideas. This package (written in C and Python) provides an `ms` compatible command line interface along with a Python API, and is freely available under the terms of the GNU GPL at https://pypi.python.org/pypi/msprime. The implementation uses a simple linked-list based representation of ancestral segments, and uses a binary indexed tree (Fenwick, 1994, 1995) to ensure the choice of ancestral segment involved in a recombination event can be done in logarithmic time. The implementation of `msprime` is based on the listings for Hudson’s algorithm given in Appendix A, which should provide sufficient detail to make implementation in a variety of languages routine.

### 2.2 Performance analysis

Surprisingly little is known about the complexity of Hudson’s algorithm. We do not know, for example, what the expected maximum number of extant ancestors is, nor the distribution of ancestral material among them. The most important unknown value in terms of quantifying the complexity of the algorithm is the expected number of events that must be generated. It is sufficient to consider the recombination events as the number of common ancestor and recombination events is approximately equal (Wiuf and Hein, 1999a). Hudson’s algorithm traverses a subset of the ARG as it generates the marginal genealogies in which we are interested, and so we know that the expected number of recombination events we encounter is less than *e^ρ^* (Ethier and Griffiths, 1990). This subset of the ARG is sometimes known as the ‘little’ ARG, but the relationship between the ‘big’ and little ARGs has not been well characterised.

Figure 1 plots the average number of recombination events generated by Hudson’s algorithm for varying sequence lengths and sample sizes. In this plot we also show the results of fitting a quadratic function to the number of recombination events as we increase the scaled recombination rate *ρ*. The fit is excellent, suggesting that the current upper bound of *e^ρ^* is far too pessimistic. Wiuf and Hein (1999a) previously noted that the observed number of events in Hudson’s algorithm was ‘subexponential’ but did not suggest a quadratic bound. Another point to note is that the rate at which the number of events grows as we increase the sample size is extremely slow, suggesting that Hudson’s algorithm should scale well for large sample sizes.

**Figure 1:**
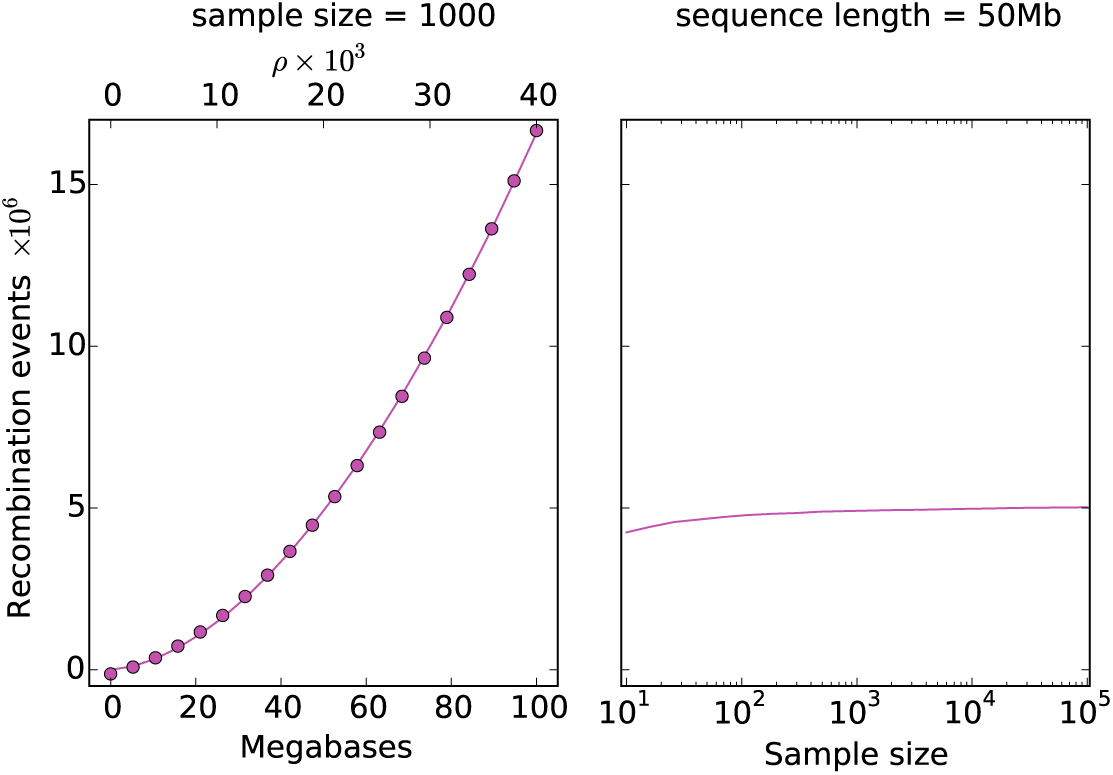
The mean number of recombination events in Hudson’s algorithm over 100 replicates for varying sequence length and sample size. In the left panel we fix *n* = 1000 and vary the sequence length. Shown in dots is a quadratic fitted to these data, which has a leading coefficient of 8.4 *×* 10^−3^. In the right panel we fix the sequence length at 50 megabases and vary the sample size.

These expectations are borne out well in observations of our implementation of Hudson’s algorithm in `msprime`. Figure 2 compares the time required to simulate coalescent trees using a number of simulation packages. As we increase the sequence length in the left-hand panel, the running time of `msprime` increases faster than linearly, but at quite a slow rate. `msprime` is faster than the SMC approximations (`MaCS` and `scrm`) until *ρ* is roughly 20000, and the difference is minor for sequence lengths greater than this. `msprime` is far faster than `msms`, the only other exact simulator in the comparison (we did not include `ms` in these comparisons as it was too slow and is unreliable for large sample sizes). As we increase the sample size in the right-hand panel, we can see that `msprime` is far faster than any other simulator. Two versions of `msprime` are shown in these plots: one outputting Newick trees (to ensure that the comparison with other simulators is fair), and another that outputs directly in `msprime`’s native format. Conversion to Newick is an expensive process, particularly for larger sample sizes. When we eliminate this bottleneck, simulation time grows at quite a slow, approximately linear rate. The memory usage of `msprime` is also modest, with the simulations in Figure 2 requiring less than a gigabyte of RAM. Supplementary Figure 8 shows that the mean number of recombination breakpoints (i.e., the number of recombination events within ancestral material) output by all these simulators is identical, and matches Hudson and Kaplan’s (1985) prediction very well, giving us some confidence in the correctness of the results. The `cosi2` (Shlyakhter et al., 2014) simulator was not included in this comparison as it does not support outputting trees directly.

**Figure 2:**
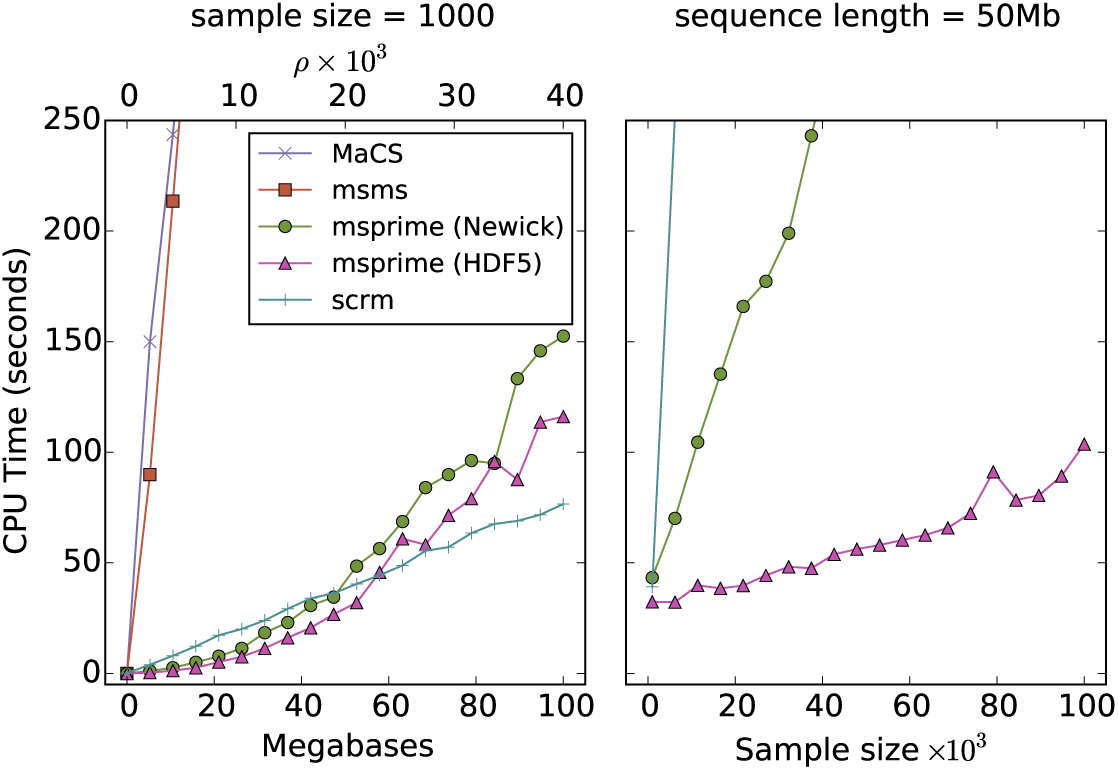
Comparison of the average running time over 100 replicates for various coalescent simulators with varying sequence length and sample size. `msms` (Ewing and Hermisson, 2010) is the most efficient published simulator based on Hudson’s algorithm that can output genealogies. `MaCS` (Chen et al., 2009) is a popular SMC based simulator, and `scrm` (Staab et al., 2014) is the most efficient sequential simulator currently available. Both `MaCS` and `scrm` were run in the most approximate SMC′ mode. Two results are shown for `msprime`; one outputting Newick trees and another outputting the native HDF5 based format.

We are often interested in the haplotypes that result from imposing a mutation process onto genealogies as well as the genealogies themselves. Supplementary Figure 6 compares the time required to generate haplotypes using `scrm`, `msprime` and `cosi2`. Simulation times are similar in all three for a fixed sample size of 1000 and increasing sequence length. For increasing sample sizes, both `cosi2` and `msprime` are substantially faster than `scrm`. However, `msprime` is significantly faster than `cosi2` (and uses less memory; see Supplementary Figure 7), particularly when we remove the large overhead of outputting the haplotypes in text form.

Performance statistics were measured on Intel Xeon E5-2680 processors running Debian 8.2. All code required to run comparisons and generate plots is available at https://github.com/jeromekelleher/msprime-paper.

## 3 Efficient genealogical analysis

There has been much recent interest in the problem of representing large scale genetic data in formats that facilitate efficient access and calculation of statistics (Durbin, 2014; Layer et al., 2015; Li, 2015). The use of ‘succinct’ data structures, which are highly compressed but also allow for efficient queries is becoming essential: the scale of the data available to researchers is so large that naive methods simply no longer work.

Although genealogies are fundamental to biology, there has been little attention to the problem of encoding trees in a form that facilitates efficient computation. The majority of research has focused on the accurate interchange of tree structures and associated metadata. The most common format for exchanging tree data is the Newick format (Felsenstein, 1989), which although ill-defined (Vos et al., 2012) has become the de-facto standard. Newick is based on the correspondence of tree structures with nested parentheses, and is a concise method of expressing tree topologies. Because of this recursive structure, specific extensions to the syntax are required to associate information with tree nodes (Maddison et al., 1997; Zmasek and Eddy, 2001). XML based formats (Han and Zmasek, 2009; Vos et al., 2012) are much more flexible, but tend to require substantially more storage space than Newick (Vos et al., 2012). Various extensions to Newick have been proposed to incorporate more general graph structures (Morin and Moret, 2006; Buendia and Narasimhan, 2006; Cardona et al., 2008; Than et al., 2008), as well as a GraphML extension to encode ARGs directly (McGill et al., 2013). Because Newick stores branch lengths rather than node times, numerical precision issues also arise when summing over many short branches (McGill et al., 2013).

**Figure 3:**
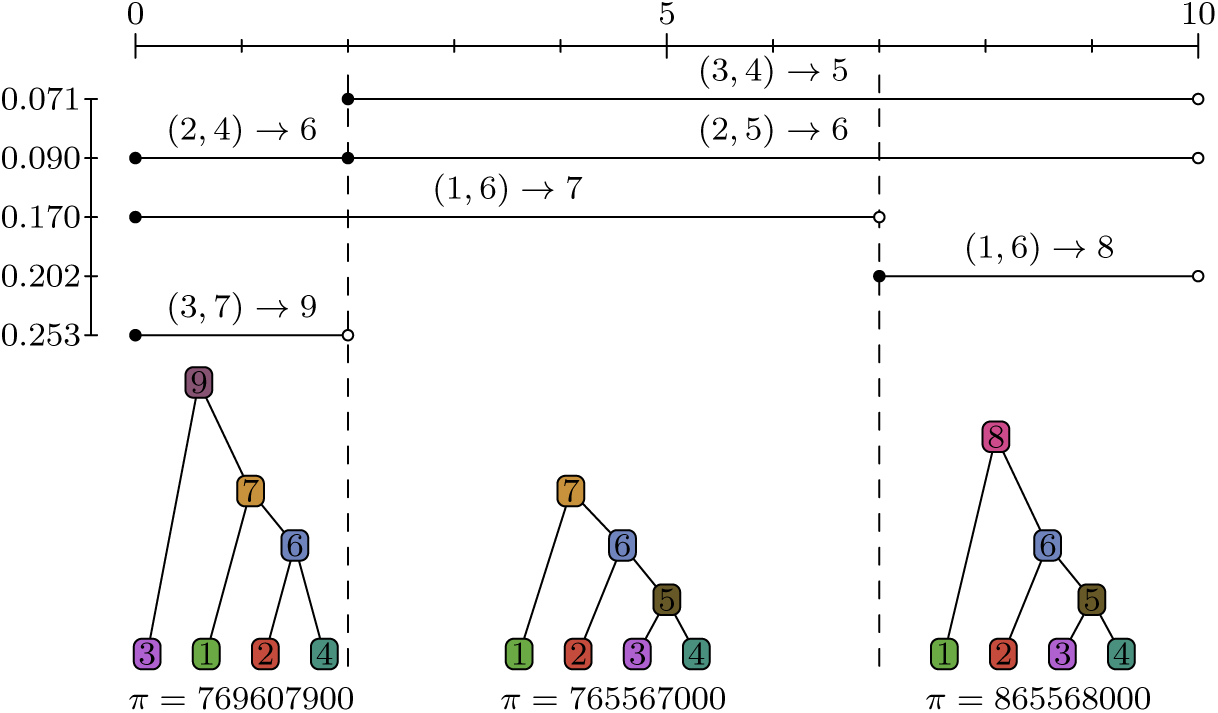
Coalescence records and corresponding marginal trees. The *x*-axis represents genomic coordinates, and *y*-axis represents time (with the present at the top). Each line segment in the top section of the figure represents a coalescence record; e.g., the first segment corresponds to the coalescence record (2, 10, 5, (3, 4), 0.071). The lower section of the figure shows the corresponding trees in pictorial and sparse tree form. We have omitted commas and brackets from this sequence representation for compactness.

General purpose Bioinformatics toolkits such as BioPerl (Stajich et al., 2002) and BioPython (Cock et al., 2009) provide basic tools to import trees in the various formats. More specific tree processing libraries such as DendroPy (Sukumaran and Holder, 2010), ETE (Huerta-Cepas et al., 2010), and APE (Paradis et al., 2004) provide more sophisticated tools such as visualisation and tree comparison algorithms. None of these libraries are designed to handle large collections of correlated trees, and cannot make use of the shared structure within a sequence of correlated genealogies. The methods employed rarely scale well to trees containing hundreds of thousands of nodes.

In this section we introduce a new representation of the correlated trees output by a coalescent simulation using coalescence records. In Section 3.1 we discuss this tree sequence structure and show how it compares in practice to existing approaches in terms of storage size. Section 3.2 presents an algorithm to sequentially generate the marginal genealogies from a tree sequence, which we compare with existing Newick-based methods. Finally, in Section 3.3 we show how the algorithm to sequentially visit trees can be extended to efficiently maintain the counts of leaves from a specific subset, and show how this can be applied in a calculation commonly used in genome wide association studies.

### 3.1 Tree Sequences

The output of Hudson’s algorithm as described in Section 2.1 is a list of coalescence records. Each coalescence record is a tuple (*ℓ, r, u, c, t*) describing the coalescence of a list of child nodes *c* into the parent *u* at time *t* over the half-closed genomic interval [*ℓ, r*). (Because only binary trees are possible in the standard coalescent, we assume the child node list *c* is a 2-tuple (*c*_1_*, c*_2_) throughout. However, arbitrary numbers of children can be accommodated without difficulty to support common ancestor events in which more than two lineages merge (Donnelly and Kurtz, 1999; Pitman, 1999; Sagitov, 1999).) We refer to this set of records as a *tree sequence*, as it is a compact encoding of the set of correlated trees representing the genealogies of a sample. Figure 3 shows an illustration of the tree sequence output by an example simulation (a full trace of this simulation is shown in Supplementary Figure 5).

The tree sequence provides a concise method of representing the correlated genealogies generated by coalescent simulations because it stores node assignments shared across adjacent trees exactly once. Consider node 7 in Figure 3. This node is shared in the first two marginal trees, and in both cases it has two children, 1 and 6. Even though the node spans two marginal trees, the node assignment is represented in one coalescence record (0, 7, 7, (1, 6), 0.170). Importantly, this holds true even though the subtree beneath 6 is different in these trees. Thus, any assignment of a pair of children to a given parent that is shared across adjacent trees will be represented by exactly one coalescence record.

Coalescence records provide a full history of the coalescence events that occurred in our simulation. (Recall that we distinguish between common ancestor events, which may or may not result in marginal coalescences, and coalescence events which are defined as a single contiguous block of genome merging within a common ancestor.) The effects of recombination events are also stored indirectly in this representation in the form of the left and right coordinate of each record. For every distinct coordinate between 0 and *m*, there must have been at least one recombination event that occurred at that breakpoint. However, there is no direct information about the times of these recombination events, and many recombinations will happen that leave no trace in the set of coalescence records. For example, if we have a recombination event that splits the ancestry of a given lineage, and this is immediately followed by a common ancestor event involving these two lineages, there will be no record of this pair of events.

On the other hand, if we consider the records in order of their left and right coordinates we can also see them as defining the way in which we transform the marginal genealogies as we move across a chromosome. Because many adjacent sites may share the same genealogy, we need only consider the coordinates of our records in order to recover the distinct genealogies and the coordinate ranges over which they are defined. To obtain the marginal tree covering the interval [0, 2), for example, we simply find all records with left coordinate equal to 0 and apply these to the empty sparse tree π. To subsequently obtain the tree corresponding to the interval [2, 7) we first remove the records that do not apply over this interval, which must have right coordinate equal to 2. In the example, this corresponds to removing the assignments (2, 4) → 6 and (3, 7) → 9. Having removed the ‘stale’ records that do not cover the current interval, we must now apply the new records that have left coordinate 2. In this case, we have two node assignments (3, 4) → 5 and (2, 5) → 6, and applying these changes to the current tree completes the transformation of the first marginal tree into the second.

There is an important point here. As we moved from left-to-right across the simulated chromosome we transitioned from one marginal tree to the next by removing and applying only two records. Crucially, modifying the nodes that were affected by this transition did not result in a relabelling of any nodes that were not affected. As Wiuf and Hein (1999a,b) showed, the effect of a recombination at a given point in the sequence is to cut the branch above some node in the tree to the left of this point, and reattach it within another branch. This process is known as a subtree-prune-and-regraft (Song, 2003, 2006) and requires a maximum of three records to express in our tree sequence formulation.

Prune-and-regraft operations that do not affect the root require three records, as illustrated in Figure 4. Two other possibilities exist for how the current tree can be edited as we move along the sequence. The first case is when we have a prune and regraft that involves a change in root node; this requires only two records and is illustrated in the first transition in Figure 3. The other case that can arise from a single recombination event is a simple root change in which the only difference between the adjacent trees is the time of the MRCA. This requires one record, and is illustrated in the second transition in Figure 3. These three possibilities are closely related to the three classes of subtree-prune-and-regraft identified by Song (2003, 2006).

**Figure 4:**
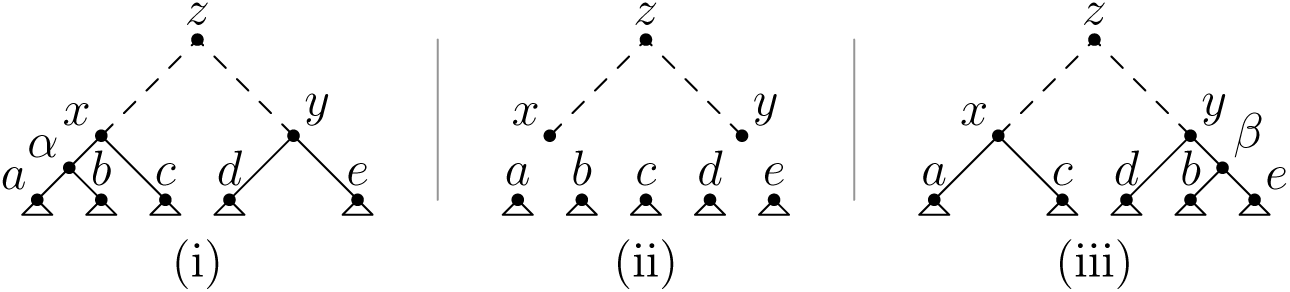
A prune and regraft not involving the root requires three records. (i) We begin with two subtrees rooted at *x* and *y*, and we wish to prune the subtree rooted at *b* and graft it in the branch joining *e* to *y*. (ii) We remove the assignments (*a*, *b*) *→ α*, (*α, c*) *→ x* and (*d, e*) → *y*. After this operation, the subtrees *a,…, e* are disconnected from the main tree. The main trunk the tree rooted at *z* is unaffected, as are the subtrees below *a,…, e*. (iii) We add the records (*a, c*) *→ x*, (*b, e*) → *β* and (*d, β*) → *y*, completing the transition.

Knowing the maximum number of records arising from a single recombination event provides us with a useful bound on the expected number of records in a tree sequence. Because the expected number of recombination events within ancestral material is approximately *ρ* log *n* (Hudson and Kaplan, 1985; Wiuf and Hein, 1999a) we know that the expected number of tree transitions is *ρ* log *n*. The number of records we require for these tree transitions is then clearly ≤ 3*p* log *n*. We also require *n* − 1 records to describe the first tree in the sequence, and so the total number of records is ≤ *n* + 3*ρ* log *n* − 1.

Storing a tree sequence as a set of coalescence records therefore requires *O*(*n* + *ρ* log *n*) space, whereas any representation that stores each tree separately (such as Newick) must require *O*(*nρ* log *n*) space. This difference is substantial in practice. As an example of a practical simulation of the sort currently being undertaken, we repeated the simulation run by Layer et al. (2015), in which we simulate a 100 megabase region with a recombination rate of 10^−3^ per base per 4*N_e_* generations for a sample of 100,000 individuals. This simulation required approximately 6 minutes and 850MB of RAM to run using `msprime`; the original simulation reportedly required over 4 weeks using `MaCS` on similar hardware.

Outputting the results as coalescence records in a simple tab-delimited text format resulted in a 173MB file (52MB when gzip compressed). In contrast, writing the trees out in Newick form required around 3.5TB of space. Because plain text is a poor format for storing structured numerical data (Kelleher et al., 2013b), `msprime` provides a tree sequence storage file format based of the HDF5 standard (The HDF Group, 1997-2015). Using this storage format, the file size is reduced to 88MB (41MB using the transparent zlib compression provided by the HDF5 library).

To compare the efficiency of storing correlated trees as coalescence records with the TreeZip compression algorithm (Matthews et al., 2010) we output the first 1000 trees in Newick format, resulting in a 3.2GB text file (1.1GB gzip compressed). The TreeZip compression algorithm required 10 hours to run and resulted in an 882MB file (83MB gzip compressed). Unfortunately, it was not feasible to run TreeZip on all 3.5TB of the Newick data, but we can see that with only around 0.1% of the input data, the compressed representation is already larger than the simple text output of the entire tree sequence when expressed as coalescence records.

Associating mutation information with a tree sequence is straightforward. For example, to represent a mutation that occurs on the branch that joins node 7 to node 9 at site 1 in Figure 3, we simply record the tuple (7, 1). (Infinite sites mutations can be readily accommodated by assuming that the coordinate space is continuous rather than discrete.) Because only the associated node and position of each mutation needs to be stored, this results in a very concise representation of the full genealogical history and mutational state of a sample. Repeating the simulation above with with a scaled mutation rate of 10^−3^ per unit of sequence length per 4*N_e_* generations resulted in 1.2 million infinite sites mutations. The total size of the HDF5 representation of the tree sequence and mutations was 102MB (49MB using HDF5’s zlib compression). In contrast, the text-based haplotype strings consumed 113GB (9.7GB gzip compressed). Converting to text haplotypes required roughly 9 minutes and 14GB of RAM.

The PBWT (Durbin, 2014) represents binary haplotype data in a format that is both highly compressed and enables efficient pattern matching algorithms. We converted the mutation data above into PBWT form, which required 22MB of storage. Thus, the PBWT is a more compact representation of a set of haplotypes than the tree sequence. However, the PBWT does not contain any genealogical data, and therefore contains less information than the tree sequence.

### 3.2 Generating trees

Coalescence records provide a very compact means of encoding correlated genealogies. Compressed representations of data usually come at the cost of increased decompression effort when we wish to access the information. In contrast, we can recover the marginal trees from a set of coalescence records orders of magnitude more quickly than is possible using existing methods. In this section we define the basic algorithm required to sequentially generate these marginal genealogies.

For algorithms involving tree sequences it is useful to regard the set of coalescence records as a table and to index the columns independently (see Supplementary Table 1 for the table corresponding to Figure 3). Therefore define a tree sequence *T* as a tuple of vectors *T* = (**l, r, u, c, t**), such that for each index 1 ≤ *j ≤ M*, (**l***_j_*, **r***_j_*, **U***_j_*, **C***_j_*, **t***_j_*) corresponds to one coalescence record output by Hudson’s algorithm, and there are *M* records in total. It is also useful to impose an ordering among the children at a node, and so we assert that ***c****_j_*_,1_ < ***c****_j_*,_2_ for all 1 ≤ *j* ≤ *M*.

If we wish to obtain the tree for a given site *x* we simply find the *n* − 1 records that intersect with this point and build the tree by applying these records. We begin by setting π*_j_ ←* 0 for 1 ≤ *j* ≤ max(**u**), and then set 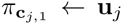 and 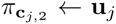 for all *j* such that l*_j_* ≤ *x* ≤ r*_j_*. Spatial indexing structures such as the segment tree (Samet, 1989) allow us to find all *k* segments out of a set of *N* that intersect with a given point in *O*(*k* + log *N*) time. Therefore, since the expected number of records is *O*(*n* + *ρ* log *n*) as shown in Section 3.1, the overall complexity of generating a single tree is *O*(*n* + log(*n* + *ρ* log *n*)).

A common requirement is to sequentially visit all trees in a tree sequence in left-to-right order. One possible way to do this would be to find all of the distinct left coordinates in the **l** vector and apply the process outlined above. However, adjacent trees are highly correlated and share much of their structure, and so this approach would be quite wasteful. A more efficient approach is given in Algorithm T below. For this algorithm we require two ‘index vectors’ 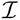 and 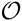 which give the indexes of the records in the order in which they are inserted and removed, respectively. Records are applied in order of nondecreasing left coordinate and increasing time, and records are removed in nondecreasing order of right coordinate and decreasing time. That is, for every pair of indexes *j* and *k* such that 1 ≤ *j* ≤ *k* ≤ *M* we have 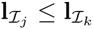 and 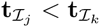, and also 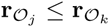 and 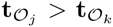. We assume that these index vectors have been pre-calculated below.

#### **Algorithm T**. (*Generate trees*).

Sequentially visit the sparse trees *π* in a tree sequence *T* = (**l, r, u, c, t**) with *M* records.

T1. [Initialisation.] Set *π_j_ ←* 0 for 1 ≤ *j ≤* max(**u**). Then set *j ←* 1, *k ←* 1 and *x* ← 0.
T2. [Insert record.] Set 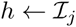, 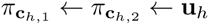, and *j ← j* + 1. If *j* ≤ *M* and 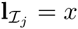, go to T2.
T3. [Visit tree.] Visit the sparse tree *π* starting at site *x*. If *j > M* terminate the algorithm. Otherwise, set 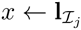.
T4. [Remove record.] Set 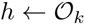, 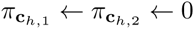 and *k ← k* + 1. Then, if 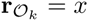 go to T4; otherwise, go to T2.

Algorithm T sequentially generates all marginal trees in a tree sequence by first applying records to the sparse tree *π* in step T2 for a given left coordinate. Once this is complete, the tree is made available to client code by ‘visiting’ it (Knuth, 2011, p.281) in T3. After the user has finished processing the current tree, we prepare to move to the next tree by removing all stale records in T4, and then return to T2. The algorithm is very efficient. Because each record is considered exactly once in step T2 and at most once in step T4 the total time required by the algorithm is *O*(*n* + *ρ* log *n*). To illustrate this efficiency, we consider the time required to iterate over the trees produced by the large example simulation used throughout this section. Reading in the full tree sequence in `msprime`’s native HDF5 based format and iterating over all 1.1 million trees using the Python API required approximately 3 seconds. In contrast, using the BioPython (Cock et al., 2009) version 1.64 Newick parser required around 3 seconds *per tree*, leading to an estimated 38 days to iterate over all trees. Similarly, ETE (Huerta-Cepas et al., 2010) version 2.3.9 required 4.5 seconds per tree, and DendroPy (Sukumaran and Holder, 2010) version 4.0.2 required around 14 seconds per tree. Comparing Python Newick parsers to `msprime` may be somewhat misleading, since the majority of `msprime`’s tree processing code is written in C. However, APE (Paradis et al., 2004) version 3.1, which uses a Newick parser written in C, also required around 7 seconds per tree. Thus, using `msprime`’s API we can iterate over this set of trees more than a *million times* faster than any of these alternatives.

Algorithm T generates only the sparse tree *π* mapping each node to its parent. It is easy to extend this algorithm to include information about the node times, children, start and end coordinates and other information. We have also assumed binary trees here, but it is trivial to extend the algorithm to work with more general trees. When computing statistics across the tree sequence it is often useful to know the specific differences between adjacent trees, as this often allows us to avoid examining the entire tree. This information is directly available in Algorithm T. The tree iteration code in `msprime`’s Python API makes all of this information available, facilitating easy tree traversal in both top-down and bottom-up fashion.

### 3.3 Counting leaves

Section 3.2 provides an algorithm to efficiently visit all marginal genealogies in a tree sequence. This algorithm can be easily augmented to maintain summaries of tree properties as we sweep across the sequence. As an example of this, we show how to augment Algorithm *T* to maintain the counts of the number of leaves from a specific set that are below each internal node. More precisely, given some subset *S* of our sample, we maintain a vector *β* such that for any node *u*, *β_u_* is the number of leaves from the set *S* below *u*. This allows us to quickly calculate allele frequencies: since each mutation is associated with a particular node *u*, *β_u_*/|S| is the frequency of the mutation within *S*. Calculating allele frequencies within specific subsets of the sample has many applications, for example calculating summary statistics such as *F_ST_* (Charlesworth and Charlesworth, 2010), and association tests in genome wide association studies (Spencer et al., 2009).

Suppose we have a tree sequence *T* and we wish to generate the sparse trees *π* as before. We now also wish to generate the vector *β*, such that *β_u_* gives the number of leaf nodes in the subtree rooted at *u* that are in the set *S* ⊆ {1,…, *n*}. We assume that the index vectors 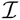 and 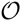 have been precomputed, as before.

#### **Algorithm L**. (*Count leaves*).

Generate the sparse trees *π* and leaf counts *β* for a tree sequence *T* = (**l, r, u, c, t**) with *M* records and set of leaves *S*.

L1. [Initialisation.] Set *π_j_ ← β_j_ ←* 0 for 1 ≤ *j ≤* max(**u**). Set *β_j_ ←* 1 for each *j* ∈ *S*. Then set *j ←* 1, *k ←* 1 and *x ←* 0.
L2. [Insert record.] Set 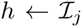, 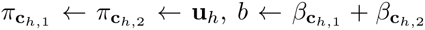 and *j ← j* + 1.
L3. [Increment leaf counts.] Set *v ←* **u***_h_*. Then, while *v* ≠ 0, set *β_v_ ← β_v_* + *b* and *v ← π_v_*. Afterwards, if *j* ≤ *M* and 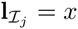, go to L2.
L4. [Visit tree.] Visit (*π, β*). If *j* > *M* terminate the algorithm; otherwise, set 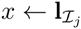.
L5. [Remove record.] Set 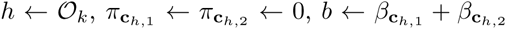 and *k ← k* + 1.
L6. [Decrement leaf counts.] Set *v →* **u***_h_*. Then, while *v* ≠ 0, set *β_v_ ← β_v_* − *b* and *v ← π_v_*. Afterwards, if 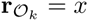, go to L5; otherwise, go to L2.

Algorithm L works in the same manner as Algorithm T: for each tree transition, we remove the stale records that no longer apply to the genomic interval currently under consideration, and apply all new records that begin at location *x*. We update the sparse tree *π* by applying a record in step L2, and then update the leaf count *β* to account for this new node assignment. In step L3 we propagate the corresponding leaf count gain up to the root, before returning to L2 if necessary. Once we have applied all of the inbound records we then visit the tree by making *π* and *β* available to the user in L4. Then, if any more trees remain, we move on by removing the outbound records in steps L5 and L6, updating *β* to account for the corresponding loss in leaf counts. The correctness of the algorithm depends on the ordering of the index vectors 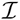 and 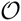. Records are always inserted in increasing order of time, and always removed in decreasing order of time within a tree transition. Therefore, for any record in which subtrees rooted at *c*_1_ and *c*_2_ become the children of *u*, we are guaranteed that these subtrees are complete and that 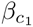 and 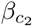 are correct. Removing outbound records in reverse order of time similarly guarantees that the leaf counts within the disconnected subtrees that we create are maintained correctly.

Algorithm L clearly examines each record at most once in steps L2 and L5. Steps L3 and L6 contain loops to propagate leaf counts up the tree, and are therefore not constant time operations. Since coalescent genealogies are asymptotically balanced (Li and Wiehe, 2013), the expected height of a tree (in terms of the number of nodes) is log_2_ *n*. Therefore, the cost of steps L3 and L6 is *O*(log_2_ *n*) per record, leading to a log_2_ *n* extra cost over Algorithm T. In practical terms, this extra cost is negligible. For example, msprime automatically maintains counts for all leaves (and optionally can maintain counts for specific subsets) when doing all tree transitions. The 3 second time quoted above required to iterate over all 1. 1 million trees in the large simulation example includes the cost of maintaining counts for all 10^5^ leaves at all internal nodes. To demonstrate this efficiency, we ran a simple genome wide association test, where we split the sample into 50,000 cases and controls. One of the most powerful and popular applications for running such association tests is `plink` (Purcell et al., 2007). After converting the simulated data to a 29G BED file, the stable version of `plink` (1.07) required 176 minutes to run a simple association test. The development version of `plink` (1.9) required 54 seconds. Using `msprime`’s Python API, the same odds-ratio test required around 10 seconds.

## 4 Discussion

The primary contribution of this paper is to introduce a new encoding for the correlated trees resulting from simulations of the coalescent with recombination. This encoding follows on from previous work in which trees are encoded as integer vectors (Kelleher et al., 2013a, 2014), but makes the crucial change that tree vectors are sparse. Using this encoding, the effects of each coalescence event are stored as simple fixed-size records that provide sufficient information to recover all marginal genealogies after the simulation has completed. This approach leads to very large gains in simulation performance over classical simulators such as `ms`, so that the exact simulation of genealogies for the coalescent with recombination over chromosome scales is feasible for the first time. We have presented an implementation based on the sparse tree encoding called `msprime`, which is faster than all other simulators for large sample sizes. Currently, we support all demographic events that can be simulated using ms in a single population. We hope to add support for simple discrete population structure in the near future, as well as for populations evolving in continuous space (Barton et al., 2010a,b, 2013b). We also hope to add gene conversion (Wiuf and Hein, 2000) and variable recombination rates following the approach taken in `cosi2`.

Coalescence records also lead to an extremely compact storage format that is several orders of magnitude smaller than the most compact method currently available. Despite this very high level of compression, accessing the genealogical data is very efficient. In an example with 100,000 samples, we saw a roughly 40,000-fold reduction in file size over the Newick tree encoding, and a greater than million-fold decrease in the time required to iterate over the genealogies compared to several popular libraries. This efficiency is gained through very simple algorithms that we have stated rigorously and unambiguously, and also analysed in terms of their computational complexity. Being able to process such large sample sizes is not an idle curiosity; on the contrary, we have a pressing need to work with such datasets. We envisage three immediate uses for our work.

Firstly, sequencing projects are being conducted on an unprecedented scale (Genome of the Netherlands Consortium et al., 2014; UK10K Consortium et al., 2015; 1000 Genomes Project Consortium et al., 2015; Gudbjartsson et al., 2015; Eisenstein, 2015; Stephens et al., 2015), and the storage and analysis of these data pose serious computational challenges. Sophisticated new methods are being developed to organise and analyse information on this immense scale (Durbin, 2014; Li, 2015; Layer et al., 2015). Perversely, developers have struggled to generate simulated data on a similar scale (Durbin, 2014; Layer et al., 2015), as present day simulators perform poorly on these huge sample sizes. Using `msprime`, the time required to generate genome scale data for hundreds of thousands of samples is reduced from weeks to minutes.

Secondly, prospective studies such as UK Biobank (Collins, 2011; Wain et al., 2015) are collecting genetic and high-dimensional phenotypic data for hundreds of thousands of samples. The key statistical method to interrogate such data is the genome wide association study (GWAS) (Manolio, 2013), and large sample size has been identified as the single most important factor in determining the power of these studies (Spencer et al., 2009). Simulation plays a key role in GWAS, and typically proceeds by superimposing the disease model of interest on haplotypes obtained via various methods (Yang et al., 2011). Because the accurate modelling of linkage disequilibrium is essential in disease genetics (Schaffner et al., 2005), recombination must be incorporated. Resampling methods (Marchini et al., 2007; Li and Li, 2008; Spencer et al., 2009; Su et al., 2011) generate simulated haplotypes based on an existing reference panel, and provide a good match to observed linkage patterns. However, there is some bias associated with this process, and there are statistical difficulties when the size of the sample required is larger than the reference panel. Other methods obtain simulated haplotypes from population genetics models via forwards-in-time (Lohmueller et al., 2008; Lohmueller, 2014) or coalescent (Günther et al., 2011; Chung and Shih, 2013) simulations. None of these methods can efficiently handle the huge sample sizes required, however. A simulator for high dimensional phenotype data based on msprime could alleviate these performance issues and be a key application for the library.

Thirdly, today’s large sample sizes provide us with an unprecedented opportunity to understand the history and geographic structure of our species. Aside from its intrinsic interest, correctly accounting for population stratification is critical for the interpretation of association studies (Marchini et al., 2004; McCarthy et al., 2008), particularly for rare variants (Mathieson and McVean, 2012, 2014). Researchers are seeking to understand fine scale population structure using methods based on principal component analysis (Novembre et al., 2008), admixture fractions (Alexander et al., 2009; Lawson et al., 2012; Liu et al., 2013), length of haplotype blocks (Ralph and Coop, 2013; Harris and Nielsen, 2013; Barton et al., 2013a) and allele frequencies (Gutenkunst et al., 2009). To date, it has been challenging to assess the accuracy of these methods, as simulations struggle to match the required sequence lengths and sample sizes. Furthermore, methods based on the SMC approximation (Li and Durbin, 2011; Schiffels and Durbin, 2014) have been tested using SMC simulations out of necessity, making it difficult to assess the impact of the approximation on accuracy. Simulations of the exact coalescent with recombination at chromosome scales for large sample sizes and arbitrary demographies will be an invaluable tool for developers of such methods.

As we have demonstrated, the tree sequence structure leads to very efficient algorithms, and allows us to encode simulated data very compactly. We would also wish to encode biological data in this structure so that we can apply these algorithms to analyse real data. However, to do this we must estimate a tree sequence from data, which is a non-trivial task. Nonetheless, there has been much work in this area (Gusfield, 2014) with several heuristic (Minichiello and Durbin, 2006) and more principled approaches that may be adopted (O’Fallon, 2013; Rasmussen et al., 2014). Using the PBWT (Durbin, 2014) to find long haplotypes (which will usually correspond to long records) seems like a particularly promising avenue.

Finally, an interesting question arises when we consider the problem of inferring a tree sequence from data. Many authors regard the ARG as a useful data structure to represent the history of a sample and explicitly infer ARGs from data (Minichiello and Durbin, 2006; Rasmussen et al., 2014). The tree sequence is not an ARG, however, since any given tree sequence can correspond to an infinite number of ARGs. By discarding information about the times and locations of recombination events, we lose information about the underlying graph that was traversed by Hudson’s algorithm. We have seen that this leads to a very compact representation and to highly efficient algorithms to reconstruct marginal trees and calculate allele frequencies. In a sense, the tree sequence is all that we can ever hope to estimate. If the mutation rate was infinite, we could reconstruct the tree sequence perfectly, but we would still not know the precise time and location of all recombination events. The specific realisation of the ARG that gave rise to the data that we observe is therefore unknowable. By focusing on the problem of estimating the observable consequences of an ARG—the tree sequence—we may increase our power to detect its properties.

## Acknowledgements

We would like to thank Richard Durbin for helpful discussions and insights. This work was supported by Wellcome Trust core award 090532/Z/09/Z to the Wellcome Trust Centre for Human Genetics, Wellcome Trust grant 100956/Z/13/Z to GM, and EPSRC grants EP/I01361X/1, EP/I013091/1 and EP/K034316/1 to AME.

## A Detailed listing for Hudson’s algorithm

In this section we provide a detailed description of our implementation of Hudson’s algorithm. First, we require some notation. Let 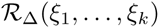 define a single independent sample from a random variable with distribution Δ and parameters ξ_1_,…,ξ*_k_*. (Note that each instance of 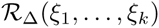 within an algorithm listing represents an *independent* random sample from the specified distribution.) Using this notation, we define 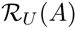 to be an element of the set *A* chosen uniformly at random, and 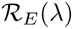 as a sample from an exponentially distributed random variable with rate λ. We use a simple linked list representation of ancestral segments such that for a segment *z*, prev(*z*) denotes the previous segment to *z* in the linked list, and similarly next(*z*) denotes the next segment. Let Λ denote a special segment indicating the end of a chain (the null reference is convenient for this purpose in many languages). Let *z* ← Segment (*ℓ, r, u, x, y*) denote a newly allocated segment such that left(*z*) = *ℓ*, right(*z*) = *r*, node(*z*) = *u*, prev(*z*) = *x* and next(*z*) = *y*. We sometimes omit the last two parameters for convenience; in this case, they are implicitly defined as Λ, and therefore Segment (*ℓ, r, u*) = Segment (*ℓ*, *r, u*, Λ, Λ). Each element of a linked list of these segments corresponds to a contiguous block of ancestry in which we map the node *u* to the half-closed interval [*ℓ*, *r*).

During recombination events we choose a breakpoint randomly and split the ancestral material within an ancestor at that point. We model these breakpoints as ‘links’ between adjacent sites. We use a binary indexed tree (Fenwick, 1994, 1995) *L* to track the cumulative number of links subtended by each extant segment (segments are ordered arbitrarily in this cumulative sum over the segments in extant ancestors). A segment *x* subtends right(*x*) − left(*x*) − 1 links if it is the first in a chain; if it is not, it subtends right(*x*) − right(prev(*x*)). That is, a segment is associated with all the links that fall both within the interval it covers and also with the links that fall in the interval between it and its predecessor. To set the number of links mapped to a segment *x* to *v*, we use the notation *L_x_* ← *v*. To find the total number of links subtended by all segments, we use total(*L*), and to obtain the cumulative number of links subtended by segment *x*, we use total(*L, x*). Finally, find(*L, v*) returns the last segment whose cumulative sum is ≤ *v*. Using these tools we can randomly choose a link and find the segment that subtends it in logarithmic time.

Termination of Hudson’s algorithm works by a gradual process of removing segments in which the MRCA has been reached. We implement this by maintaining a map *S* that counts the number of extant segments intersecting with a given interval. We use a balanced binary tree (Knuth, 1998, §6.2.3) to store this map. To assign a value *v* to key *k*, we write *S_k_* ← *v*. The data structure supports two further operations: search(*S, k*) returns the largest key ≤ *k*, and nextkey (*S, k*) returns the smallest key > *k*. For each key *k*, *S_k_* counts the number of extant segments in the interval [*k*, nextkey(*S*, *k*)). As the simulation proceeds we update this map to account for coalescences that occur, inserting keys and decrementing the counts as necessary.

### **Algorithm H**. (*Hudson’s algorithm*).

Simulate the coalescent with recombination for a sample of *n* individuals on a sequence of *m* sites with recombination at rate *r* per generation between adjacent sites.

H1. [Initialisation.] Set 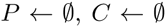, *S* ← BalancedBinaryTree() and *L* ← BinaryIndexedTree(). Then, for 1 ≤ *j* ≤ *n*, set *x* ← Segment(0, *m, j*), *L_x_* ← *m* − 1 and *P* ← *P* ⋃ {*x*}. Finally, set *S*_0_ ← *n*, *S_m_* ← −1, *w* ← *n* +1 and *t* ← 0.
H2. [Event.] Set λ*_r_* ← *r* total(*L*), *λ* ← λ*_r_* + |*P*| (|*P*| − 1), and set 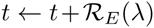. If 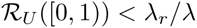, invoke Algorithm R; otherwise, invoke Algorithm C.
H3. [Loop.] If |*P*| ≠ 0 go to H2.

The basic structure of Hudson’s algorithm is very simple. We begin in H1 by allocating the set *P* to represent the extant ancestors and *C* to store our coalescence records. We also allocate the balanced binary tree *S* and the binary indexed tree *L* as discussed above. We then allocate a segment *x* covering the interval [0, *m*), that points to node *j* for each individual 1 ≤ *j* ≤ *n* in the sample, record that this segment subtends *m* − 1 links and then insert it into the set of ancestors *P*. Afterwards, we initialise the map *S* by setting *S*_0_ ← *n* and *S_m_* ← −1 (stating that the number of extant segments in the interval [0, *m*) is *n*), set the next available node *w* to *n* + 1 and our clock *t* to zero.

In H2, we calculate the current rate of recombination and common ancestor events, and increment *t* accordingly. We then choose the type of the next event and invoke either Algorithm *R* or Algorithm C. Once the appropriate subroutine has completed, we move on to H3, where we either terminate or loop back to H2. Upon termination, *C* contains the set of coalescence records that defines the output of the algorithm.

Algorithm *R* implements a single recombination event by choosing a link uniformly and breaking it, resulting in a new individual being added to the set of extant ancestors. There are two possibilities for this link: it is either between two segments or within a segment, and these possibilities are dealt with separately in steps R2 and R3, respectively. In either case, *z* points to the head of the segment chain representing the new individual, which is inserted into *P* in step R4.

### **Algorithm R**. (*Recombination event*).

Choose a link uniformly and break it, resulting in one extra individual in the set of extant ancestors.

R1. [Choose link.] Set 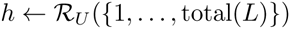, *y* ← find(*L, h*), *k* ← right(*y*) − total(*L, y*) + *h* − 1 and *x* ← prev(*y*). Then, if left(*y*) < *k* go to R3.
R2. [Break between segments.] Set next(*x*) ← Λ, prev(*y*) ← Λ, *z* ← *y* and go to R4.
R3. [Break within segment.] Set *z* ← Segment(*k*, right(*y*), node(*y*), Λ, next(*y*)). Then, if next(*y*) ≠ Λ, set prev(next(*y*)) ← *z*. Afterwards, set next(*y*) ← Λ, right(*y*) ← *k* and *L_y_* ← *L_y_* + *k* − right(*z*).
R4. [Update population] Set *L_z_* ← right(*z*) − left(*z*) − 1 and *P* ← *P* ⋃ {*z*}.

The algorithm begins in step *R*1 by choosing a link *h* uniformly from the total(*L*) that are currently being tracked. We then find the segment *y* that subtends this link using the binary indexed tree `find` function. Once we have found the segment in question, we then calculate the corresponding breakpoint *k*, so that we can determine whether link *h* falls within *y* or between *y* and its predecessor *x*. Thus, if the breakpoint *k* > left(*y*), we go to R3, and otherwise proceed to step R2.

Step R2 is very straightforward. Because the breakpoint *k* is between the two segments *x* and *y*, we must simply break the forward and reverse links in the segment chain between them. After breaking these links, we now have an independent segment chain starting with *z*, which represents the new individual to be added to the set of ancestors. On the other hand, if the breakpoint *k* falls within *y*, we must split this segment in step R3 such that the ancestral material from left(*y*) to *k* remains assigned to the current individual and the remainder is assigned to the new individual *z*. We must also update the number of links subtended by the segment *y*, which has right(*z*) – *k* fewer links as a result of this operation. Finally, step R4 inserts the segment *z* into the set of ancestors, since this is the first segment in the new individual. However, we must also update the information about the number of links subtended by this segment. Since *z* is the head of a new segment chain, there is no previous segment, and the number of links it subtends is right(*z*) − left(*z*) − 1. After this, we complete the recombination event, returning to Algorithm H.

Algorithm C implements a single common ancestor event, where we choose two individuals randomly and merge their ancestral segment chains. If these two ancestors have overlapping segments we record the corresponding coalescence events. When a coalescence occurs, we decrement the number of extant segments in the corresponding interval by updating *S*. When this value is reduced to 1, we discard the corresponding segment since it can have no further effect on the genealogies we are interested in. Thus, the algorithm always removes two individuals from the set of ancestors *P*, but may reinsert zero or one, depending on whether any ancestral segments remain after merging. By this process the size of *P* is eventually reduced to zero and Hudson’s algorithm is complete.

### **Algorithm C**. (*Common ancestor event*).

Choose two ancestors uniformly and merge their segments, recording any coalescences that occur as a consequence.

C1. [Choose ancestors.] Set 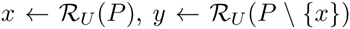. Then, set *P* ← *P* \ {*x, y*}, *z* ← Λ and *c* ← 0.
C2. [Loop head] If *x* = Λ and *y* = Λ, terminate the algorithm. Set α ← Λ. If *x* ≠ Λ and *y* ≠ Λ go to C3. Otherwise, if *x* ≠ A set α ← *x* and set *x* ← Λ. If *y* ≠ Λ set α ← *y* and set *y* ← Λ. Go to C8.
C3. [Choose case] If left(*y*) < left(*x*), set *β* ← *x*, *x* ← *y* and *y* ← *β*. Then, if right(*x*) ≤ left(*y*), set α ← *x*, *x* ← next(*x*), next(*α*) ← Λ and go to C8; otherwise, if left(*x*) ≠ left(*y*) set α ← Segment(left(*x*), left(*y*), node(*x*)), left(*x*) ← left(*y*) and go to C8.
C4. [Coalescence] If *c* = 0, set *c* ← 1 and *w* ← *w* + 1. Afterwards, set *u* ← *w* − 1, *ℓ* ← left(*x*) and *r\** ← min(right(*x*), right(*y*)). If *ℓ ∉ S*, set *j* ← search(*S, ℓ*) and *S_ℓ_* ← *S_j_*. Similarly, if r* *∉ S*, set *j* ← search(*S, r\**) and 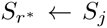. Then, if *S_l_* ≠ 2 go to C6.
C5. [Segment MRCA] Set *S_ℓ_* ← 0 and *r* ← nextkey(*S, ℓ*). Go to C7.
C6. [Decrement overlaps.] Set *r* ← *ℓ*. Then, while *S_r_* ≠ 2 and *r* < *r*\*, set *S_r_* ← *S_r_* − 1 and *r* ← nextkey(*S, r*). Afterwards, set α ← Segment(*ℓ, r, u*).
C7. [Update *x* and *y*] Set *C* ← *C* ⋃ {(*ℓ, r*, node(*x*), node(*y*), *u, t*)}. If right(*x*) = *r*, set *x* ← next(*x*); otherwise, set left(*x*) ← *r*. If right(*y*) = *r*, set *y* ← next(*y*); otherwise, set left(*y*) ← *r*.
C8. [Update links] If α = Λ go to C2. If *z* = Λ set *P* ← *P* ⋃ {*α*} and *L_α_* ← right(*α*) − left(*α*) − 1; otherwise, set next(*z*) ← *α* and *L_α_* ← right(*α*) − right(*z*). Afterwards, set prev(*α*) ← *z*, *z* ← *α* and go to C2

We begin in step C1 by choosing our individuals *x* and *y* and removing them from the set of ancestors. We then set the tail of the segment chain representing the common ancestor *z* to the null segment Λ, and then proceed into the main loop of the algorithm. This loop is controlled in step C2, and works by taking the leading segment from the *x* and *y* chains at each iteration and processing it. Once all segments have been consumed, we exit. Therefore, if both *x* and *y* are null, this loop has completed and we terminate the algorithm. Otherwise, we set *α* to the null segment. Throughout, we use this variable to point to the next segment that is to be merged into the segment chain representing the ancestor of the two chosen individuals. The last-merged segment in this chain is pointed to by *z*, and the necessary operations to include α into the global state are carried out in step C8.

Returning to the head of the loop in C2, if either *x* or *y* is null we have reached the end of one of the segment chains, and all that remains to do is attach the remainder of the non-null chain to our new individual. If both *x* and *y* are non-null, on the other hand, we proceed to C3. In this step we consider the two segments *x* and *y* and decide which of a number of cases we must deal with. First, we maintain the invariant that left(*x*) ≤ right(*y*); if this is violated, we swap the variables. Then, we address the various cases that can occur as *x* and *y* overlap.

The simplest case is when there is no overlap between *x* and *y* which occurs when right(*x*) ≤ left(*y*); here, we simply merge *x* into the new segment chain and move on to C8. The next case we deal with is when we have a partial overlap between *x* and *y*, which occurs when left(*x*) ≠ right(*y*). In this case, we create a new segment to represent this ‘overhang’, and merge this into the new segment chain in C8. Finally, if none of these conditions have been satisfied, we know that left(*x*) = right(*y*) and there is therefore a coalescence which we handle in C4.

First, we check if another coalescence has occurred during this common ancestor event. If not, we set our flag *c* ← 1, and increment the next node w. Afterwards, we set the parent node for this coalescence *u*, and set *ℓ* and *r*\* to the boundaries of the coalescing interval. We then check if *ℓ* and *r*\* are in *S* so that we can subsequently update the number of extant segments in the intervals to account for the coalescence. There are then two possibilities: if *S*_ℓ_ = 2, we know that the MRCA has been reached in an interval starting at *ℓ*, which we deal with in C5; if not, we move on to C6.

In general, there will be many intervals with different numbers of extant segments overlapping between *ℓ* and *r*\*. In C6 we iterate over each of these intervals, decrementing the number of extant segments to account for the current coalescence. After this has completed, we allocate the new segment *α* and move on to C7. Here, we record the coalescence by updating the set C, handle any trailing overlaps that may occur, and update *x* and *y* to point to the appropriate next segments in their respective chains.

Step C8 is the final step of the main loop, where we insert the new segment *α* into the chain representing the common ancestor. Firstly, if this segment is null as a result of reaching the MRCA, then we have nothing to do, and so return to the start of the main loop. The variable *z* is used to keep track of the previous segment that was merged into the common ancestor’s segment chain. Thus, if *z* is null we know that *α* is the first segment in the new chain and so we can use this opportunity to insert the new individual into the set of ancestors *P*; otherwise, we merge *α* into the existing chain. In both cases, we update the number of links subtended by *α* as appropriate, before returning to C2.

As stated, Algorithm H correctly simulates the coalescent with recombination and returns a set of coalescence records fully describing the generated genealogies. In the interest of brevity we have omitted some details that are important for efficiency. Firstly, it is important to defragment segments in order to save time and memory. That is, if we have two adjacent segments (*ℓ, k, u*) and (*k, r, u*) we should merge these into a single equivalent segment (*ℓ, r, u*). This can be done quite simply after Algorithm C has completed, and we can detect when such defragmentation is required in step C8. Similarly, it is vital for efficiency to opportunistically defragment the map that counts the number of extant segments in a given interval. Since *S_j_* counts the number of segments covering the interval [*j, k*), where *k* is the smallest key > *j* in *S*, if *S_j_* = *S_k_* we can simply delete the key *k* without loss of information. Although it does not affect simulation efficiency, it is also important to defragment the coalescence records output by the algorithm. This is easily done, since any records (*ℓ, k, u, c, t*) and (*k, r, u, c, t*) that can be merged must be stored sequentially without any intervening records.

The implementation of `msprime` is closely based on Algorithm `H` as given here. We also provide a simpler Python implementation in the file `algorithms.py` at https://github.com/jeromekelleher/msprime-paper. This repository also contains all code required to run the simulations, and to create all figures and illustrations in this paper.

## B Illustration of Hudson’s algorithm

Figure 5 shows an illustration of Hudson’s algorithm for a sample of four individuals. In this illustration we show the state of the algorithm and its effects on the marginal trees after every event. The state of the algorithm is fully defined by the ancestral lineages (defined by the segments of ancestral material that they carry), the next available node *w* and the current time *t*. Although it is not necessary to store the partially built genealogies in memory, we show them here in the lower part of each panel for clarity. The left-to-right axis represents genomic coordinates. We also show the current time (*t*) and the number of potential recombination breakpoints (*L*) in each panel.

**Figure 5:**
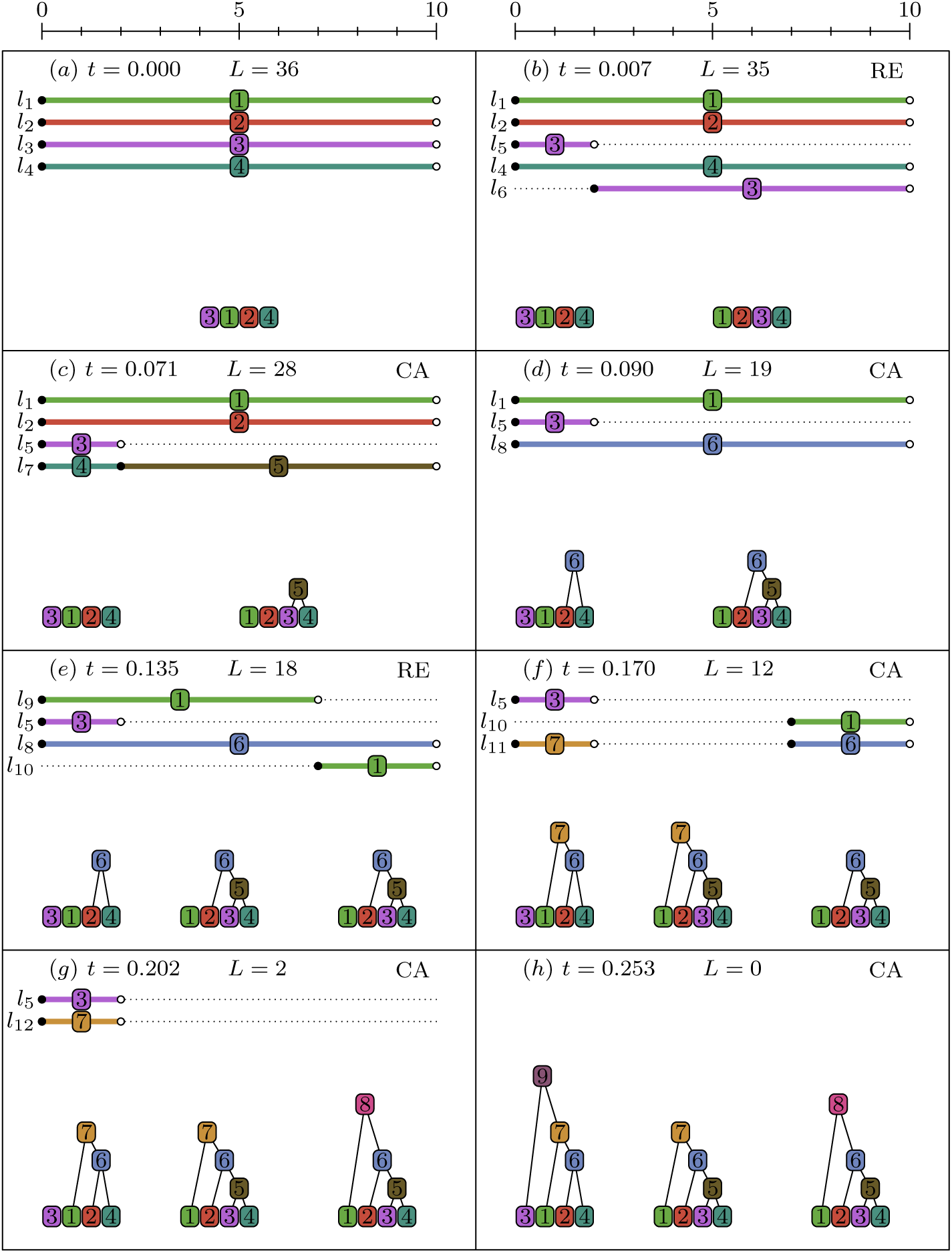
An illustration of Hudson’s algorithm using sparse trees. In each panel we show the state of the algorithm after an event. Events are either recombination (RE) or common ancestor (CA). On the top of each panel, every line represents an ancestor which may be composed of several distinct segments. The bottom of each panel shows the state of the trees at that point in time. The horizontal direction represents genomic coordinates.

In this example, we have simulated the ancestry of the sample for a sequence of 10 sites. The initial state of the simulation at time 0 is shown in panel (a), where we see four lineages corresponding to our sampled chromosomes. Lineage *l*_1_ can be represented as the segment (0, 10, 1), which states that over the genomic interval [0, 10), the lineage occupies the tree node 1. This information is shown explicitly in the figure, where we draw the full range of each segment and label the line with the node it is associated with. Nodes are colour-coded, so that we can easily see which tree nodes are associated with each segment. Since this is the initial state of the algorithm, the only tree nodes defined are the leaf nodes. This is shown in the bottom part of the panel, where we draw out the nodes of the trees that have been assigned so far. (The nodes are ordered in these panels such that they are consistent with the orderings induced by later events.)

The first event that we encounter as we go backwards in time is a recombination event which occurs at time *t* = 0.007. Panel (b) shows the state of the simulation immediately after this event. Recombination has split lineage *l*_3_, resulting in two new lineages, *l*_5_ and *l*_6_. As the breakpoint was at 2, we have *l*_5_ = (0, 2, 3) and *l_6_* = (2, 10, 3). The other effect of this recombination event is to create a new tree: since the histories of the sample over the intervals [0, 2) and [2, 10) can now be different, we must create a new tree to record these histories as they are simulated. (Note again that these trees are shown for illustration only; they are not stored in the simulation.)

After the recombination in event (b), a common ancestor event occurs in (c) in which *l*_4_ and *l*_6_ are merged to form a common ancestor *l*_7_. At a common ancestor event we merge the ancestral material from two lineages. Any nonoverlapping segments are copied directly into the new lineage. In this example, only one of the lineages carried ancestral material in the interval [0, 2), and so this is copied directly to the common ancestor. However, in the interval [2, 10) both carry ancestral material, and so a coalescence occurs. In this coalescence, nodes 3 and 4 have a common ancestor in the interval [2, 10). We therefore create a new node 5, and update the tree covering the interval [2, 10) to reflect this.

The simulation continues generating common ancestor and recombination events at the relevant rates until complete genealogies have been generated across the entire sequence. Termination of the algorithm is controlled by keeping track of the amount of ancestral material present in each distinct interval produced by recombination. An important aspect of Hudson’s algorithm is that we do not continue to track the ancestry of segments in which the trees are already complete. An example of this can be seen in panel (f) of Figure 5. In this event lineages *l*_8_ and *l*_9_ have merged to form *l*_11_. In the interval [0, 7), these have overlapping ancestral material and we therefore create a new node 7 and update the trees covering [0, 2) and [2, 7) to show that node 7 is the parent of 1 and 6. However, we note that the tree covering the interval [2, 7) is complete as a result, and so we omit the segment mapping to the new node over this interval. This process is important for efficiency, as we would continue to generate recombination and common ancestor events for the segment, even though these events could not effect the genealogy over this interval.

Panel (f) also illustrates the concept of trapped material. Lineage *l*_11_ consists of the two segments (0, 2, 7) and (7, 10, 6). Recombination events occurring anywhere in [0, 10) on this lineage will therefore result in a different arrangement of ancestral material. The total number of possible recombination breakpoints for *l*_11_ is therefore 9. In contrast, there are only 2 possible breakpoints for *l*_10_, since any recombination that occurs in [0, 7) cannot affect the ancestral material. Similarly *l*_5_ has only one potential breakpoint, and so the total number of potential breakpoints *L* = 12.

## C Supplemental Figures

**Table 1:**
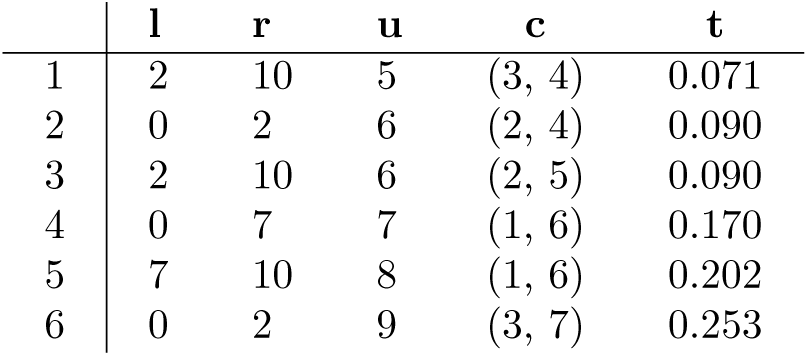
Tabular representation of the coalescence records output by the simulation in Figure 5 and depicted in Figure 3. The corresponding index vectors are 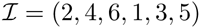 and 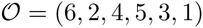.

**Figure 6:**
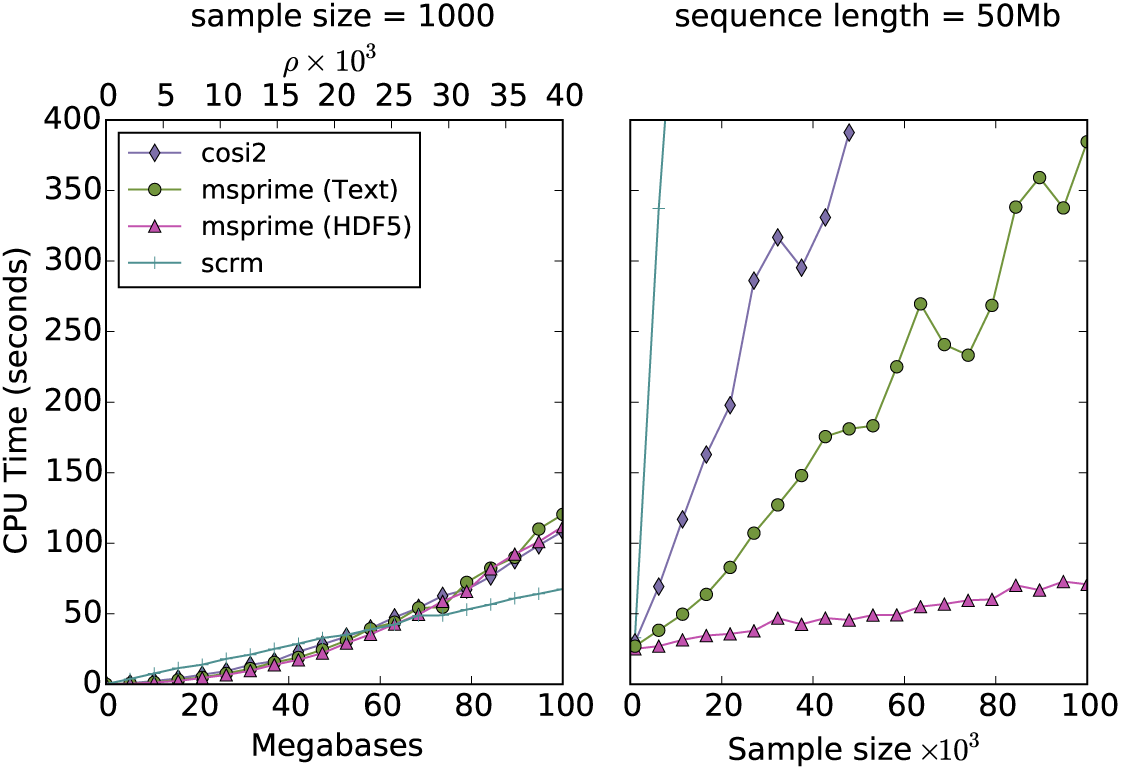
Comparisons of the running times for various coalescent simulators to generate mutations for varying sequence length and sample size. We use a scaled mutation rate of *θ* = 4*Ν_e_μ* = 0.0004.

**Figure 7:**
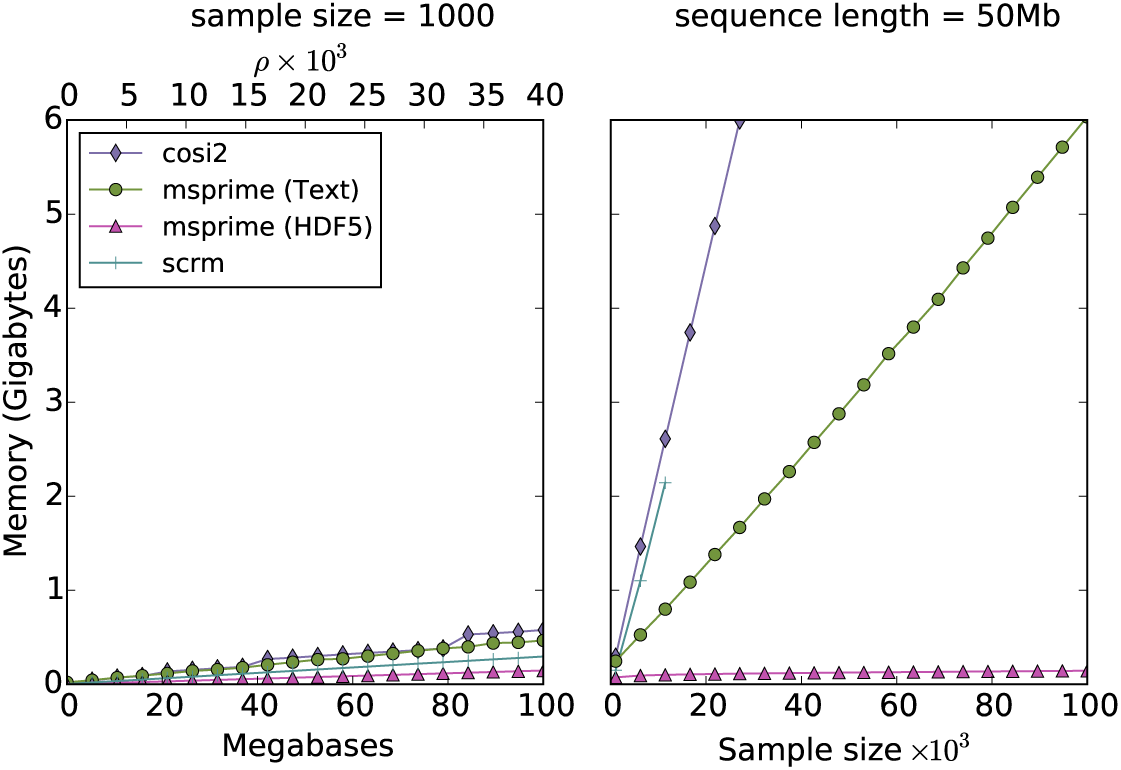
The corresponding maximum memory usages for the simulators in Figure 6.

**Figure 8:**
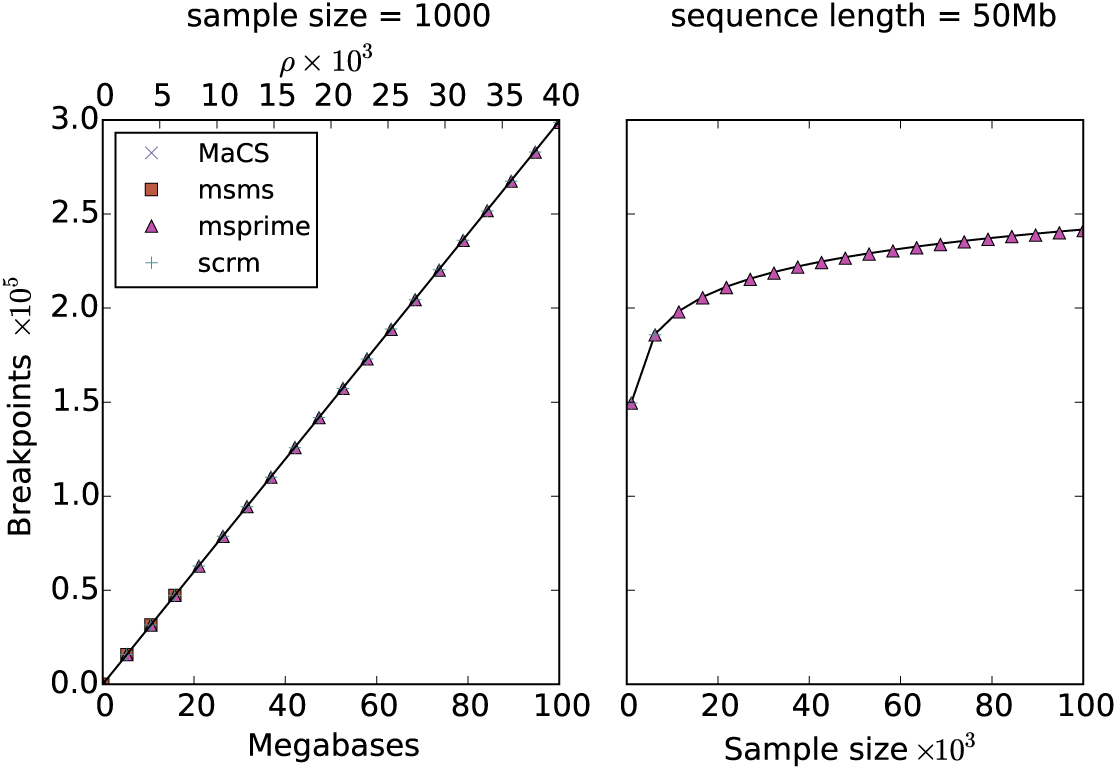
The mean number of recombination breakpoints for the simulations in Figure 2 along with the theoretical prediction (black line). This plot shows that the number of recombination events within ancestral material for these simulations is identical for all simulators and agrees very well with the theoretical value of *ρH_n_* _− 1_, where *H_n_* is the *n*th Harmonic number.

## References

1000 Genomes Project Consortium et al. A global reference for human genetic variation. Nature, 526(7571):68–74, 2015.

D. H. Alexander, J. Novembre, and K. Lange. Fast model-based estimation of ancestry in unrelated individuals. Genome Res, 19(9):1655–1664, 2009.

C. N. Anderson, U. Ramakrishnan, Y. L. Chan, and E. A. Hadly. Serial simcoal: a population genetics model for data from multiple populations and points in time. Bioinformatics, 21(8):1733–1734, 2005.

M. Arenas. Simulation of molecular data under diverse evolutionary scenarios. PLoS Comput Biol, 8(5):e1002495, 2012.

M. Arenas and D. Posada. Recodon: coalescent simulation of coding DNA sequences with recombination, migration and demography. BMC Bioinformatics, 8(1):458, 2007.

M. Arenas and D. Posada. Coalescent simulation of intracodon recombination. Genetics, 184(2):429–437, 2010.

N. H. Barton, A. M. Etheridge, and A. Véber. A new model for evolution in a spatial continuum. Electron J of Probab, 15:7, 2010a.

N. H. Barton, J. Kelleher, and A. M. Etheridge. A new model for extinction and recolonisation in two dimensions: quantifying phylogeography. Evolution, 64(9):2701–2715, 2010b.

N. H. Barton, A. M. Etheridge, J. Kelleher, and A. Véber. Inference in two dimensions: allele frequencies versus lengths of shared sequence blocks. Theor Popul Biol, 87:105–119, 2013a.

N. H. Barton, A. M. Etheridge, and A. Véber. Modelling evolution in a spatial continuum. J Stat Mech, P01002, 2013b.

A. Bhaskar, A. G. Clark, and Y. S. Song. Distortion of genealogical properties when the sample is very large. Proc Natl Acad Sci USA, 111(6):2385–2390, 2014.

P. Buendia and G. Narasimhan. Serial NetEvolve: a flexible utility for generating serially-sampled sequences along a tree or recombinant network. Bioinformatics, 22(18):2313–2314, 2006.

G. Cardona, F. Rosselló, and G. Valiente. Extended Newick: it is time for a standard representation of phylogenetic networks. BMC Bioinformatics, 9:532, 2008.

A. Carvajal-Rodríguez. Simulation of genomes: a review. Curr Genomics, 9(3):155, 2008.

B. Charlesworth and D. Charlesworth. Elements of Evolutionary Genetics. Roberts and Company, Greenwood Village, Colorado, 2010.

G. K. Chen, P. Marjoram, and J. D. Wall. Fast and flexible simulation of DNA sequence data. Genome Res, 19:136–142, 2009.

R.-H. Chung and C.-C. Shih. SeqSIMLA: a sequence and phenotype simulation tool for complex disease studies. BMC Bioinformatics, 14(1):199, 2013.

P. J. A. Cock, T. Antao, J. T. Chang, B. A. Chapman, C. J. Cox, A. Dalke, I. Friedberg, T. Hamelryck, F. Kauff, B. Wilczynski, and M. J. L. de Hoon. Biopython: freely available Python tools for computational molecular biology and bioinformatics. Bioinformatics, 25(11):1422–1423, 2009.

R. Collins. UK biobank: the need for large prospective epidemiological studies. J Epidemiol Community Health, 65(1):A37, 2011.

P. Donnelly and T. G. Kurtz. Particle representations for measure-valued population models. Ann Probab, 27(1):166–205, 1999.

R. Durbin. Efficient haplotype matching and storage using the positional Burrows-Wheeler transform (PBWT). Bioinformatics, 30(9):1266–1272, 2014.

M. Eisenstein. Big data: The power of petabytes. Nature, 527(7576):S2–S4, 2015.

A. Eriksson, B. Mahjani, and B. Mehlig. Sequential Markov coalescent algorithms for population models with demographic structure. Theor Popul Biol, 76(2):84–91, 2009.

S. N. Ethier and R. C. Griffiths. On the two-locus sampling distribution. J Math Biol, 29:131–159, 1990.

G. Ewing and J. Hermisson. MSMS: a coalescent simulation program including recombination, demographic structure, and selection at a single locus. Bioinformatics, 26(16):2064–2065, 2010.

L. Excoffier and M. Foll. fastsimcoal: a continuous-time coalescent simulator of genomic diversity under arbitrarily complex evolutionary scenarios. Bioinformatics, 27(9):1332–1334, 2011.

L. Excoffier, J. Novembre, and S. Schneider. SIMCOAL: a general coalescent program for the simulation of molecular data in interconnected populations with arbitrary demography. J Hered, 91(6):506–509, 2000.

J. Felsenstein. PHYLIP—phylogeny inference package (version 3.2). Cladistics, 5:164–166, 1989.

P. M. Fenwick. A new data structure for cumulative frequency tables. Software: Practice and Experience, 24:327–336, 1994.

P. M. Fenwick. A new data structure for cumulative frequency tables: an improved frequency-to-symbol algorithm. Technical Report 110, The University of Auckland, Department of Computer Science, 1995.

Genome of the Netherlands Consortium et al. Whole-genome sequence variation, population structure and demographic history of the Dutch population. Nat Genet, 46(8):818–825, 2014.

R. C. Griffiths. The two-locus ancestral graph. Lecture Notes-Monograph Series, 18:100–117, 1991.

R. C. Griffiths and P. Marjoram. An ancestral recombination graph. In P. Donnelly and S. Tavaré, editors, Progress in Population Genetics and Human Evolution, IMA Volumes in Mathematics and its Applications, volume 87, pages 257–270. Springer-Verlag, Berlin, 1997.

D. F. Gudbjartsson, H. Helgason, S. A. Gudjonsson, F. Zink, A. Oddson, A. Gylfason, S. Besenbacher, G. Magnusson, B. V. Halldorsson, E. Hjartarson, et al. Large-scale whole-genome sequencing of the Icelandic population. Nat Genet, 47(5):435–444, 2015.

T. Gunther, I. Gawenda, and K. J. Schmid. phenosim - a software to simulate phenotypes for testing in genome-wide association studies. BMC Bioinformatics, 12(1):265, 2011.

D. Gusfield. ReCombinatorics. MIT Press, Cambridge Massachusetts, 2014.

R. N. Gutenkunst, R. D. Hernandez, S. H. Williamson, and C. D. Bustamante. Inferring the joint demographic history of multiple populations from multidimensional SNP frequency data. PLoS Genet, 5(10):e1000695, 2009.

M. V. Han and C. M. Zmasek. phyloXML: XML for evolutionary biology and comparative genomics. BMC Bioinformatics, 10(356), 2009.

K. Harris and R. Nielsen. Inferring demographic history from a spectrum of shared haplotype lengths. PLoS Genet, 9(6):e1003521, 2013.

G. Hellenthal and M. Stephens. mshot: modifying Hudson’s ms simulator to incorporate crossover and gene conversion hotspots. Bioinformatics, 23(4):520–521, 2007.

S. Hoban, G. Bertorelle, and O. E. Gaggiotti. Computer simulations: tools for population and evolutionary genetics. Nat Rev Genet, 13(2):110–122, 2012.

R. R. Hudson. Testing the constant-rate neutral allele model with protein se quence data. Evolution, 37(1):203–217, 1983a.

R. R. Hudson. Properties of a neutral allele model with intragenic recombination. Theor Popul Biol, 23:183–201, 1983b.

R. R. Hudson. Gene genealogies and the coalescent process. Oxford Surveys in Evolutionary Biology, 7:1–44, 1990.

R. R. Hudson. Generating samples under a Wright-Fisher neutral model of genetic variation. Bioinformatics, 18(2):337–338, 2002.

R. R. Hudson and N. Kaplan. Statistical properties of the number of recombination events in the history of a sample of DNA sequences. Genetics, 111(1):147–164, 1985.

J. Huerta-Cepas, J. Dopazo, and T. Gabaldón. ETE: a python environment for tree exploration. BMC Bioinformatics, 11:24, 2010.

N. Kaplan and R. R. Hudson. The use of sample genealogies for studying a selectively neutral m-loci model with recombination. Theor Popul Biol, 28:382–396, 1985.

J. Kelleher, N. H. Barton, and A. M. Etheridge. Coalescent simulation in continuous space. Bioinformatics, 29(7):955–956, 2013a.

J. Kelleher, R. W. Ness, and D. L. Halligan. Processing genome scale tabular data with wormtable. BMC Bioinformatics, 14:356, 2013b.

J. Kelleher, A. M. Etheridge, and N. H. Barton. Coalecent simulation in continuous space: algorithms for large neighbourhood size. Theor Popul Biol, 95:13–23, 2014.

J. F. C. Kingman. The coalescent. Stoch. Proc. Appl., 13(3):235–248, 1982.

D. E. Knuth. Sorting and Searching, volume 3 of The Art of Computer Programming. Addison-Wesley, Reading, Massachusetts, second edition, 1998.

D. E. Knuth. Combinatorial Algorithms, Part 1, volume 4A of The Art of Computer Programming. Addison-Wesley, Upper Saddle River, New Jersey, 2011.

G. Laval and L. Excoffier. SIMCOAL 2.0: a program to simulate genomic diversity over large recombining regions in a subdivided population with a complex history. Bioinformatics, 20(15):2485–2487, 2004.

D. J. Lawson, G. Hellenthal, S. Myers, and D. Falush. Inference of population structure using dense haplotype data. PLoS Genet, 8(1):e1002453–e1002453, 2012.

R. M. Layer, N. Kindlon, K. J. Karczewski, Exome Aggregation Consortium, and A. R. Quinlan. Efficient genotype compression and analysis of large genetic-variation data sets. Nat Methods, 2015. doi: doi:10.1038/nmeth.3654.

C. Li and M. Li. GWAsimulator: a rapid whole-genome simulation program. Bioinformatics, 24(1):140–142, 2008.

H. Li. BGT: efficient and flexible genotype query across many samples. Bioinformatics, 2015. doi: 10.1093/bioinformatics/btv613.

H. Li and R. Durbin. Inference of human population history from individual whole-genome sequences. Nature, 475:493–496, 2011.

H. Li and T. Wiehe. Coalescent tree imbalance and a simple test for selective sweeps based on microsatellite variation. PLoS Comput Biol, 9(5):e1003060, 2013.

L. Liang, S. Zöllner, and G. R. Abecasis. GENOME: a rapid coalescent-based whole genome simulator. Bioinformatics, 23(12):1565–1567, 2007.

M. Liang and R. Nielsen. The lengths of admixture tracts. Genetics, 197:953–967, 2014.

Y. Liu, G. Athanasiadis, and M. E. Weale. A survey of genetic simulation software for population and epidemiological studies. Hum Genomics, 3(1):79, 2008.

Y. Liu, T. Nyunoya, S. Leng, S. A. Belinsky, Y. Tesfaigzi, and S. Bruse. Softwares and methods for estimating genetic ancestry in human populations. Hum Genomics, 7(1), 2013.

K. E. Lohmueller. The impact of population demography and selection on the genetic architecture of complex traits. PLoS Genet, 10(5):e1004379, 2014.

K. E. Lohmueller, A. R. Indap, S. Schmidt, A. R. Boyko, R. D. Hernandez, M. J. Hubisz, J. J. Sninsky, T. J. White, S. R. Sunyaev, R. Nielsen, et al. Proportionally more deleterious genetic variation in European than in African populations. Nature, 451(7181):994–997, 2008.

D. R. Maddison, D. L. Swofford, and W. P. Maddison. Nexus: An extensible file format for systematic information. Syst Biol, 46(4):590–621, 1997.

T. Mailund, M. H. Schierup, C. N. Pedersen, P. J. Mechlenborg, J. N. Madsen, and L. Schauser. CoaSim: a flexible environment for simulating genetic data under coalescent models. BMC Bioinformatics, 6(1):252, 2005.

T. A. Manolio. Bringing genome-wide association findings into clinical use. Nat Rev Genet, 14(8):549–558, 2013.

J. Marchini, L. R. Cardon, M. S. Phillips, and P. Donnelly. The effects of human population structure on large genetic association studies. Nat Genet, 36(5):512–517, 2004.

J. Marchini, B. Howie, S. Myers, G. McVean, and P. Donnelly. A new multipoint method for genome-wide association studies by imputation of genotypes. Nat Genet, 39(7):906–913, 2007.

P. Marjoram and J. D. Wall. Fast “coalescent” simulation. BMC Genet, 7:16, 2006.

I. Mathieson and G. McVean. Differential confounding of rare and common variants in spatially structured populations. Nat Genet, 44(3):243–246, 2012.

I. Mathieson and G. McVean. Demography and the age of rare variants. PLoS Genet, 10(8):e1004528, 2014.

S. J. Matthews, S.-J. Sul, and T. L. Williams. A novel approach for compressing phylogenetic trees. In M. Borodovsky, J. Gogarten, T. Przytycka, and S. Rajasekaran, editors, Bioinformatics Research and Applications, volume 6053 of Lecture Notes in Computer Science, pages 113–124. Springer Berlin Heidelberg, 2010.

M. I. McCarthy, G. R. Abecasis, L. R. Cardon, D. B. Goldstein, J. Little, J. P. Ioannidis, and J. N. Hirschhorn. Genome-wide association studies for complex traits: consensus, uncertainty and challenges. Nat Rev Genet, 9(5):356–369, 2008.

J. R. McGill, E. A. Walkup, and M. K. Kuhner. GraphML specializations to codify ancestral recombinant graphs. Fron Genet, 4:146, 2013.

G. A. T. McVean and N. J. Cardin. Approximating the coalescent with recombination. Philos Trans R Soc Lond B Biol Sci, 360:1387–1393, 2005.

M. J. Minichiello and R. Durbin. Mapping trait loci by use of inferred ancestral recombination graphs. Am J Hum Genet, 79:910–922, 2006.

M. M. Morin and B. M. E. Moret. NetGen: generating phylogenetic networks with diploid hybrids. Bioinformatics, 22(15):1921–1923, 2006.

J. Novembre, T. Johnson, K. Bryc, Z. Kutalik, A. R. Boyko, A. Auton, A. Indap, K. S. King, S. Bergmann, M. R. Nelson, et al. Genes mirror geography within Europe. Nature, 456(7218):98–101, 2008.

B. D. O’Fallon. ACG: rapid inference of population history from recombining nucleotide sequences. BMC Bioinformatics, 14(1):40, 2013.

E. Paradis, J. Claude, and K. Strimmer. APE: analyses of phylogenetics and evolution in *R* language. Bioinformatics, 20:289–290, 2004.

S. Peischl, E. Koch, R. Guerrero, and M. Kirkpatrick. A sequential coalescent algorithm for chromosomal inversions. Heredity, 111:200–209, 2013.

J. Pitman. Coalescents with multiple collisions. Ann Probab, 27(4):1870–1902, 1999.

S. Purcell, B. Neale, K. Todd-Brown, L. Thomas, M. A. Ferreira, D. Bender, J. Maller, P. Sklar, P. I. de Bakker, M. J. Daly, and P. C. Sham. PLINK: a tool set for whole-genome association and population-based linkage analyses. Am J Hum Genet, 81(3):559–575, 2007.

P. Ralph and G. Coop. The geography of recent genetic ancestry across Europe. PLoS Biol, 11(5):e1001555, 2013.

S. E. Ramos-Onsins and T. Mitchell-Olds. Mlcoalsim: multilocus coalescent simulations. Evol Bioinform Online, 3:41, 2007.

M. D. Rasmussen, M. J. Hubisz, I. Gronau, and A. Siepel. Genome-wide inference of ancestral recombination graphs. PLoS Genet, 10(5):e1004342, 2014.

S. Sagitov. The general coalescent with asynchronous mergers of ancestral lines. J Appl Probab, 36(4):1116–1125, 1999.

H. Samet. The Design and Analysis of Spatial Data Structures. Addison-Wesley, Upper Saddle River, New Jersey, 1989.

S. F. Schaffner, C. Foo, S. Gabriel, D. Reich, M. J. Daly, and D. Altshuler. Calibrating a coalescent simulation of human genome sequence variation. Genome Res, 15(11):1576–1583, 2005.

S. Schiffels and R. Durbin. Inferring human population size and separation history from multiple genome sequences. Nat Genet, 46:919–925, 2014.

I. Shlyakhter, P. C. Sabeti, and S. F. Schaffner. Cosi2: an efficient simulator of exact and approximate coalescent with selection. Bioinformatics, 30(23):3427–3429, 2014.

Y. S. Song. On the combinatorics of rooted binary phylogenetic trees. Ann Comb, 7(3):365–379, 2003.

Y. S. Song. Properties of subtree-prune-and-regraft operations on totally-ordered phylogenetic trees. Ann Comb, 10(1):147–163, 2006.

C. C. Spencer and G. Coop. SelSim: a program to simulate population genetic data with natural selection and recombination. Bioinformatics, 20(18):3673–3675, 2004.

C. C. Spencer, Z. Su, P. Donnelly, and J. Marchini. Designing genome-wide association studies: sample size, power, imputation, and the choice of genotyping chip. PLoS Genet, 5(5):e1000477, 2009.

P. R. Staab, S. Zhu, D. Metzler, and G. Lunter. scrm: efficiently simulating long sequences using the approximated coalescent with recombination. Bioinformatics, 31(10):1680–1682, 2014.

J. E. Stajich, D. Block, K. Boulez, S. E. Brenner, S. A. Chervitz, C. Dagdigian, G. Fuellen, J. G. Gilbert, I. Korf, H. Lapp, H. Lehväslaiho, C. Matsalla, C. J. Mungall, B. I. Osborne, M. R. Pocock, P. Schattner, M. Senger, L. D. Stein, E. Stupka, M. D. Wilkinson, and E. Birney. The Bioperl toolkit: Perl modules for the life sciences. Genome Res, 12(10):1611–1618, 2002.

Z. D. Stephens, S. Y. Lee, F. Faghri, R. H. Campbell, C. Zhai, M. J. Efron, R. Iyer, M. C. Schatz, S. Sinha, and G. E. Robinson. Big data: Astronomical or genomical? PLoS Biol, 13(7):e1002195, 2015.

Z. Su, J. Marchini, and P. Donnelly. HAPGEN2: simulation of multiple disease SNPs. Bioinformatics, 27(16):2304–2305, 2011.

J. Sukumaran and M. T. Holder. DendroPy: a Python library for phylogenetic computing. Bioinformatics, 26(12):1569–1571, 2010.

K. M. Teshima and H. Innan. mbs: modifying hudson’s ms software to generate samples of DNA sequences with a biallelic site under selection. BMC Bioinformatics, 10(1):166, 2009.

C. Than, D. Ruths, and L. Nakhleh. PhyloNet: a software package for analyzing and reconstructing reticulate evolutionary relationships. BMC Bioinformatics, 9:322, 2008.

The HDF Group. Hierarchical Data Format, version 5, 1997–2015. http://www.hdfgroup.org/HDF5/.

UK10K Consortium et al. The UK10K project identifies rare variants in health and disease. Nature, 526(7571):82–90, 2015.

R. A. Vos, J. P. Balhoff, J. A. Caravas, M. T. Holder, H. Lapp, W. P. Maddison, P. E. Midford, A. Priyam, J. Sukumaran, X. Xia, and A. Stoltzfus. NeXML: rich, extensible, and verifiable representation of comparative data and metadata. Syst Biol, 61(4):675–689, 2012.

L. V. Wain, N. Shrine, S. Miller, V. E. Jackson, I. Ntalla, M. S. Artigas, C. K. Billington, A. K. Kheirallah, R. Allen, J. P. Cook, et al. Novel insights into the genetics of smoking behaviour, lung function, and chronic obstructive pulmonary disease (UK BiLEVE): a genetic association study in UK Biobank. Lancet Respir Med, 3(10):769–781, 2015.

J. Wakeley. Coalescent theory: an introduction. Roberts and Company, Englewood, Colorado, 2008.

J. Wakeley and T. Takahashi. Gene genealogies when the sample size exceeds the effective size of the population. Mol Biol Evol, 20(2):208–213, 2003.

Y. Wang, Y. Zhou, L. Li, X. Chen, Y. Liu, Z.-M. Ma, and S. Xu. A new method for modeling coalescent processes with recombination. BMC Bioinformatics, 15(1):273, 2014.

C. Wiuf and J. Hein. Recombination as a point process along sequences. Theor Popul Biol, 55(3):248–259, 1999a.

C. Wiuf and J. Hein. The ancestry of a sample of sequences subject to recombination. Genetics, 151(3):1217–1228, 1999b.

C. Wiuf and J. Hein. The coalescent with gene conversion. Genetics, 155(1):451–462, 2000.

J. Yang, S. H. Lee, M. E. Goddard, and P. M. Visscher. GCTA: a tool for genome-wide complex trait analysis. Am J Hum Genet, 88(1):76–82, 2011.

T. Yang, H.-W. Deng, and T. Niu. Critical assessment of coalescent simulators in modeling recombination hotspots in genomic sequences. BMC Bioinformatics, 15:3, 2014.

X. Yuan, D. J. Miller, J. Zhang, D. Herrington, and Y. Wang. An overview of population genetic data simulation. J Comput Biol, 19(1):42–54, 2012.

S. Zhu, J. H. Degnan, S. J. Goldstien, and B. Eldon. Hybrid-Lambda: simulation of multiple merger and Kingman gene genealogies in species networks and species trees. BMC Bioinformatics, 16(292), 2015.

C. M. Zmasek and S. R. Eddy. ATV: display and manipulation of annotated phylogenetic trees. Bioinformatics, 17(4):383–384, 2001.

